# Compressed representations underpin knowledge awareness in sequence learning

**DOI:** 10.64898/2026.08.22.746376

**Authors:** Haoran Jiang, Xiuyan Guo, Yang Lu, Shaobo Xu, Guanglong Liu, Xianting Shen, Xue Weng

## Abstract

Humans can acquire sequence structure before its presence enters awareness, and in some learners this implicit knowledge later becomes accessible and reportable. Yet the neural reformatting that enables this transition remains unclear. Here we used electro/magnetoencephalography (EEG/MEG) to resolve how category-based triplet rules were organized before and after awareness emerged. We found that implicit sequence knowledge was already represented in an abstract, compressed format. Within this format, rule information generalized across triplet elements, and regular-sequence elements became more similar while perceptual-category structure remained separable. Compression rate quantified how efficiently local rule information entered this shared format, and its pre-awareness level predicted later awareness specifically in the event-related potential P3 window. MEG source imaging localised pretransition compression to left middle temporal cortex and showed post-awareness expansion into prefrontal cortex. Pre-awareness directional connectivity further showed that temporal-to-prefrontal theta-band transfer was rule-specific and predicted later awareness. Overall, these findings identify a mechanism supporting conscious access to abstract sequence-rule knowledge, involving cross-element representational compression and theta-band routing from the compression source to prefrontal cortex.

## Main

Humans can learn regularities before they can report them [1]. This dissociation raises a central question for learning and consciousness: what change in neural representation makes latent structure consciously accessible [2, 3]? Sequence learning provides a tractable model for this question. Learners can extract temporal regularities from repeated experience, yet only some later become aware of the underlying rule [4–11]. This implicit-to-explicit transition is not merely an improvement in behavioural performance, but a change in access: learned structure becomes available for report, flexible use and cognitive control [12]. The representational mechanisms that prepare this transition remain poorly understood.

One possibility is that awareness emerges when sequence information becomes sufficiently strong. Yet strength alone may be insufficient [13]. For learned structure to become flexibly accessible, it may also need to be represented in a format that generalizes beyond the specific items and positions through which it was acquired [14, 15]. Such abstraction is central to flexible cognition: abstract representations support rule use across contexts [16], and sequence representations can evolve from perceptual coding of serial items toward higher-order structure that transcends sensory features [17, 18]. It remains unknown, however, whether this abstract format is already present before awareness, whether it predicts later awareness emergence and how it is integrated across cortical systems.

We treated abstraction as representational compression across sequence elements. A local sequence code can distinguish regular from control contexts within a given element or position, but such local discriminability does not establish that the rule is represented abstractly [19, 20]. An abstract sequence code should generalize across elements [14, 19, 21, 22]: a classifier trained to distinguish regular from control sequences using one element should transfer to another element with a different perceptual category and ordinal position. We quantified this cross-element generalization using cross-condition generalization performance (CCGP) based on multivariate pattern analysis (MVPA) [23]. Above-chance CCGP indicates that different elements embedded in the same regular structure share a common neural feature space [24]. We refer to this shared structure as representational compression because it reduces element-specific variability while preserving information about sequence regularity.

Representational compression provides a way to test whether abstract sequence knowledge is a precursor to awareness rather than merely its consequence. Conscious abstract knowledge is unlikely to depend only on stronger local evidence; it should also be organized in a format that bridges sensory representational contexts and supports flexible use beyond the elements through which it was acquired [21]. On this view, awareness should emerge not when sequence evidence simply accumulates, but when that evidence is expressed in a shared, element-general format. Because compression can be measured during implicit learning, before verbal report, it allows us to ask whether such a format is already in place before awareness and whether it predicts later transition. To dissociate cross-element abstraction from local decodability, we complemented CCGP with a compression rate that normalizes cross-element decoding by matched within-element decoding, estimating the proportion of locally decodable sequence information expressed in a shared format.

If compression prepares conscious access, it should also engage network-level mechanisms that make local representations available for report. Global workspace accounts propose that conscious access depends on the propagation of information from posterior representational systems to higher-order cortical regions, rather than on local encoding alone [18, 25]. This predicts that pre-transition compression should be detectable in regions encoding the relevant sequence structure and should interact with regions implicated in flexible access and report. Source-resolved magnetoencephalography (MEG) allowed us to localise cortical compression and test whether directed transfer from the compression source predicted later awareness emergence.

Here, we tested this account across three experiments combining sequence-learning tasks with EEG and MEG. Learners viewed object triplets governed by a hidden category-level sequence rule, with awareness assessed by verbal reports, reaction-time switchpoints and postlearning awareness tests under varying report and motor demands. Across experiments, sequence regularity was encoded in a compressed neural format that generalized across elements before learners could report the rule. This predictive compression was aligned with a P3 time window, localised by MEG source imaging to the middle temporal gyrus and linked to theta-band directed transfer from temporal to prefrontal cortex. These findings suggest that explicit sequence knowledge is prepared by compression within posterior representational systems and supported by theta-band communication to frontal access systems.

## Results

### Study design

We used a common sequence-rule learning framework across experiments (Fig. 1a). Learners viewed triplets composed of three object categories (face, tool, animal). Regular triplets followed a fixed order with a face in Position 1 (Type I: face–tool–animal; Type II: face–animal–tool; counterbalanced across learners), whereas control triplets violated the rule and excluded face-leading orders, ensuring separability at the triplet level and enabling element-wise generalization tests (Supplementary Table 1).

**Fig. 1:**
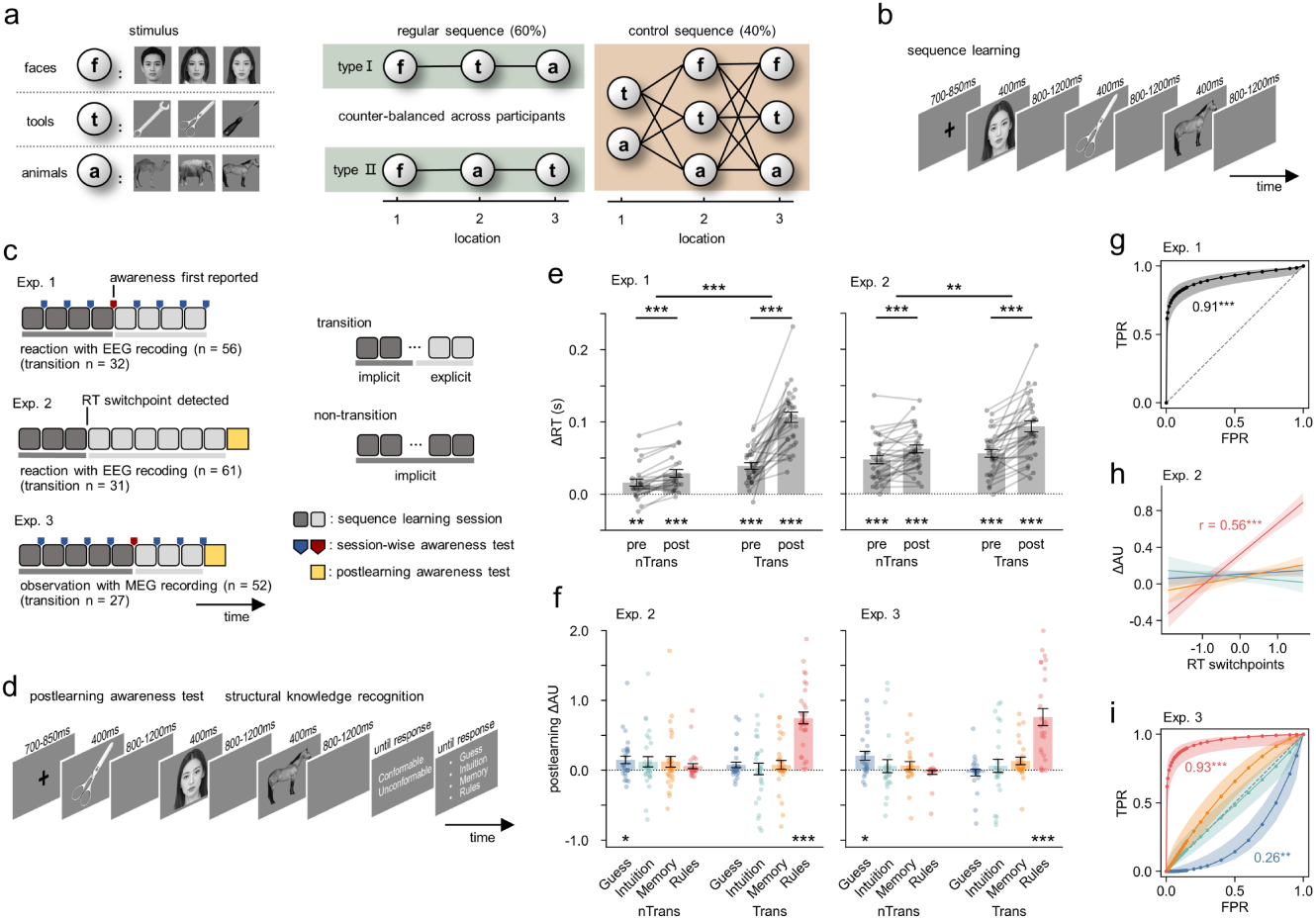
Study design and behavioural results. **a**, Triplets used in Exps. 1–3. Each category (face, tool, animal) contained three exemplars randomly sampled on each presentation. **b**, Example trial structure for a regular triplet. **c**, Overview of Exps. 1–3, including session structure, transition definition, and neural recording modality. In Exp. 2, the transition session was defined by the best-fitting switchpoint model on the ΔRT trajectory (Methods; individual estimates in Supplementary Table 3). **d**, Postlearning awareness test in Exps. 2 and 3. Learners judged whether a triplet conformed to the trained rule and reported the basis of each judgement (Guess, Intuition, Memory, or Rules; Methods). **e**, RT learning effects in Exps. 1 and 2. ΔRT = RT_control_ − RT_regular_. For transition learners (Trans), pre-to-post ΔRT increased from 39 ± 25 ms (mean ± SD) to 106 ± 40 ms in Exp. 1, and from 56 ± 30 ms to 93 ± 42 ms in Exp. 2; for non-transition learners (nTrans), it increased from 16 ± 24 ms to 29 ± 26 ms in Exp. 1, and from 47 ± 31 ms to 62 ± 30 ms in Exp. 2. Bars indicate condition means, dots show individual learners, and error bars denote s.e.m. Asterisks indicate significance (\**P <* 0.05, \*\**P <* 0.01, \*\*\**P <* 0.001). **f**, Postlearning structural knowledge in Exps. 2 and 3 quantified by rule-attributed ΔAU. Plot conventions as in **e**. **g**, Fitted ROC analysis of RT switchpoint evidence predicting report-based transition status in Exp. 1. Shading around fitted ROC curves denotes pointwise s.e. **h**, Association between RT switchpoint evidence and rule-attributed ΔAU in Exp. 2 (Pearson correlation; line shows linear fit with s.e.m.). **i**, ROC analysis of attribution-specific ΔAU predicting report-based transition status in Exp. 3. AUCs [95% CIs]: Guess, 0.26 [0.12, 0.41] (*P*_two-sided_ = 0.004; below chance); Intuition, 0.53 [0.37, 0.69]; Memory, 0.60 [0.44, 0.75] (*P*s _two-sided_ ≥ 0.211 for Intuition and Memory); Rule, 0.93 [0.86, 0.99] (*P*_one-sided_ *<* 0.001). All faces shown in this figure are fully synthetic, AI-generated images and do not depict any real individual. No photographs of participants, authors, or other identifiable persons were used as inputs or references.

Each triplet constituted one trial (Fig. 1b). Regular and control trials were interleaved in an event-related design (Extended Data Fig. 1a). Trial order was pseudorandomised to maintain stable exposure across learning, keeping the cumulative regular:control count near 3:2 from trial 30 onward (Extended Data Fig. 1b).

Task demands and awareness assessment varied across experiments (Fig. 1c) to test robustness to report reactivity and motor demands. Exp. 1 used a three-choice SRT task (8 sessions; 30 regular and 20 control per session) with session-wise subjective reports to localise awareness emergence (Supplementary Fig. 1). Exp. 2 used a no-report stream (9 sessions; 27 regular and 18 control per session) and indexed emergence via an individual RT switchpoint (Methods). Exp. 3 adopted an object-viewing paradigm with the Exp. 1 session structure and a session-wise awareness test. In Exps. 2–3, a postlearning awareness test provided an independent index of explicit knowledge (Fig. 1d).

Learners were labelled transition or non-transition, with transition-session position defined by session-wise verbal reports (Exps. 1 and 3) or an RT switchpoint (Exp. 2; Methods). Because cross-element decoding requires estimating representations within each state under category- and position-matching, learners with fewer than three sessions in either the pre- or post-transition state were excluded to ensure sufficient trials for reliable within-state decoding. Non-transition learners were assigned pseudo-transition sessions by sampling the empirical transition-position distribution to align trajectories across groups (Methods).

Neural recordings targeted cross-element generalization of sequence knowledge. Exps. 1–2 used 128-channel EEG, and Exp. 3 used 306-channel MEG, leveraging its sensitivity to fast representational dynamics in the absence of overt responses, consistent with prior work on knowledge replay [26].

### Awareness and learning effects

Despite the varied task demands, transition rates were comparable across experiments (0.57, 0.51, and 0.52 for Exps. 1–3; pairwise chi-square tests, *χ*²(1) ≤ 0.47, *P*s ≥ 0.493, *ϕ*s ≤ 0.07; Extended Data Fig. 1c). Both transition and non-transition learners in Exps. 1 and 2 showed a reliable online learning effect, quantified as ΔRT (RT_control_ − RT_regular_), across states (one-sample *t* tests against zero, *t*s ≥ 8.42, *P*s *<* 0.001, *d_z_*s ≥ 0.67; Fig. 1e; see Extended Data Fig. 2a for session-wise RT effects). Transition learners additionally showed a behavioural gain associated with awareness emergence, evidenced by a larger pre-to-post increase in ΔRT than non-transition learners (group × state interaction: *F*s ≥ 9.11, *P*s ≤ 0.004, 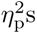 ≥ 0.13; Fig. 1e), who served as a control group to rule out time-dependent changes unrelated to transition. Switch-point modelling of the ΔRT trajectory over sessions in transition learners localised the behavioural gain to a discrete switchpoint at the transition session (Methods; Extended Data Fig. 2b–c).

The postlearning test in Exps. 2 and 3 assessed explicit knowledge by requiring learners to discriminate the trained regular sequence from an alternative sequence counterbalanced across participants (type II for type I-trained learners, and vice versa) and from control sequences (Supplementary Table 2). Attribution utility (AU) quantified each attribution category’s contribution to above-chance discrimination by jointly capturing attribution-specific accuracy, defined as the percent correct among trials assigned to that attribution (Supplementary Fig. 2a), and attribution proportion, defined as the fraction of all correct responses accounted for by that attribution (Supplementary Fig. 2b). We computed AU separately for the trained and alternative regular sequences and used their difference (ΔAU = AU_trained_ − AU_alternative_) as the structural learning index, controlling for test-induced learning (Methods). Consistent with the transition labeling, transition learners showed robust rule-attributed (explicit) ΔAU (0.75 ± 0.46 and 0.76 ± 0.63 in Exps. 2 and 3, respectively; one-sample *t* test against zero, *t*s ≥ 6.26, Bonferroni-corrected *P*s *<* 0.001, *d_z_*s ≥ 1.20), whereas non-transition learners showed modest ΔAU when discrimination was attributed to Guess (implicit; 0.15 ± 0.29 and 0.21 ± 0.32 in Exps. 2 and 3, respectively; one-sample *t*s ≥ 2.78, Bonferroni-corrected *P*s ≤ 0.038, *d_z_*s ≥ 0.50; Fig. 1f).

We evaluated pairwise consistency among the operational indices of awareness used across experiments. In Exp. 1, verbal reports and RT switchpoints showed strong agreement in classifying transition status, as assessed using ROC analysis (AUC [95% CI] = 0.91 [0.82, 0.98], *P*_one-sided_ *<* 0.001; Fig. 1g). In Exp. 2, RT switchpoints were positively associated with postlearning rule-attributed ΔAU (Pearson *r* = 0.56 [0.36, 0.71], *P <* 0.001; Fig. 1h). In Exp. 3, verbal reports and postlearning rule-attributed ΔAU also showed strong agreement (AUC = 0.93 [0.86, 0.99], *P*_one-sided_ *<* 0.001; Fig. 1i).

### Compressive knowledge representation

We tested whether sequence regularity is expressed in a compressive code, such that different triplet elements share a common neural feature space. Cross-element generalization was quantified with MVPA as cross-condition generalization performance (CCGP; Methods) using neural data with minimal motor contamination (motor-evoked EEG components removed in Exps. 1–2; Methods; Extended Data Fig. 3a; no overt responses in Exp. 3; Methods). For each participant, logistic classifiers were trained to discriminate regular from control trials on element-specific epochs (−0.2 to 1.2 s) from Positions 2–3 matched for category and ordinal position, and tested on held-out epochs from the remaining positions, including Position 1 (Fig. 2a). Ordinal-position effects in regular sequences were minimized by counterbalancing element positions across learners (tool and animal swapped between Positions 2 and 3 across Types I and II). Aggregating transfer performance across train–test configurations yielded CCGP; above-chance values indicate a shared feature space for sequence regularity across elements, isolating sequence-structure representations from perceptual, positional, and motor confounds. CCGP revealed significant sequence compression, reflected by above-chance time windows identified with a 1,000-permutation cluster-corrected test (*P*s *<* 0.05; Exp. 1: 0.25–1.01 s; Exp. 2: 0.15–1.06 s; Exp. 3: 0.32–1.06 s; Fig. 2b; Extended Data Fig. 3b). At the participant level, mean CCGP across learning positively correlated with the mean ΔRT in Exps. 1 and 2 (*r*s ≥ 0.47, *P*s ≤ 0.001) and with overall postlearning discrimination accuracy in Exps. 2 and 3 (*r*s ≥ 0.28, *P*s ≤ 0.041; Supplementary Fig. 3), linking neural compression to behavioural knowledge expression. Transition learners exhibited a larger pre-to-post increase in CCGP than non-transition learners (group × state interaction: *F*s ≥ 4.24, *P*s ≤ 0.045, 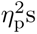 ≥ 0.08; Fig. 2c). These results suggest that compressive sequence representations can emerge prior to explicit awareness but are strengthened when the learned regularity becomes consciously accessible.

**Fig. 2:**
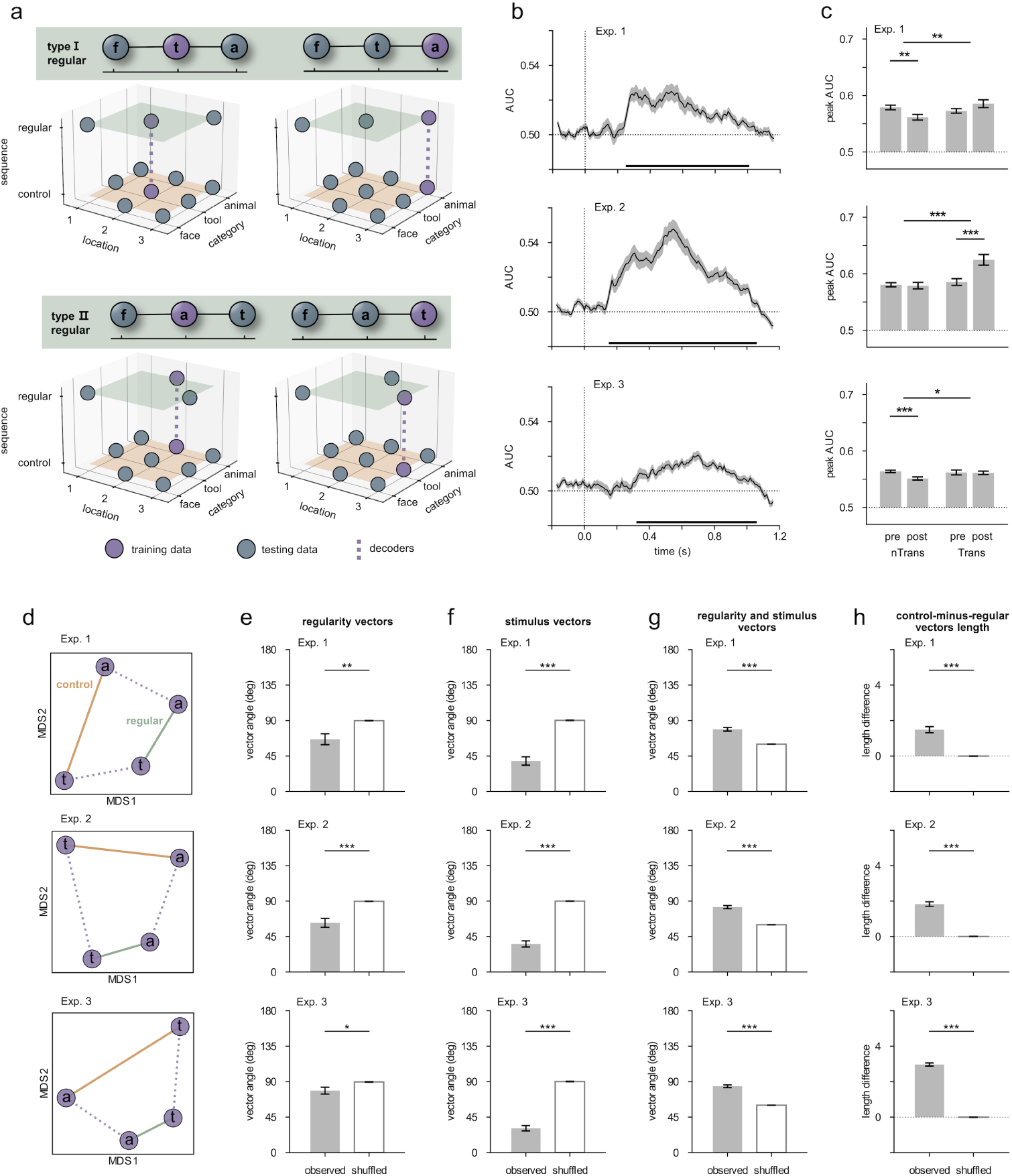
Compressive representations of sequence knowledge. **a**, Cross-element decoding. Purple and blue points indicate training and testing conditions, respectively. Dashed line denotes the decoder axis. **b**, Time-resolved CCGP (AUC) averaged across train–test configurations. Time 0 marks stimulus onset; shading denotes s.e.m.; horizontal bars mark cluster-corrected above-chance intervals. **c**, Peak CCGP across learning states for transition and non-transition groups in Exps. 1–3. **d**, MDS embedding of matched regular and control elements illustrating stimulus vectors (within sequence type) and regularity vectors (control→regular for matched elements). **e–g**, Vector geometry: alignment of regularity vectors (**e**), alignment of stimulus vectors (**f**), and separation between regularity and stimulus axes (**g**). **h**, Compactness: control–regular differences in within-sequence distances (positive values indicate tighter clustering for regular elements). Statistics were assessed against shuffled controls using two-tailed paired *t* tests.

We unpacked the cross-element compression implied by CCGP using representational dissimilarity matrices (RDMs) and multidimensional scaling (MDS). Both analyses were performed on the same element-specific epochs within the CCGP-defined significant time window, yielding condition-averaged patterns for the 2 × 2 design (tool vs. animal) × (regular vs. control). Pairwise distances between these patterns formed a 4×4 RDM (Extended Data Fig. 4), which showed reduced tool–animal dissimilarity within regular sequences relative to control sequences (*t*s ≥ 8.66, *P*s *<* 0.001, *d_z_*s ≥ 1.15), indicating increased representational overlap under the learned regularity.

The RDMs were embedded into a two-dimensional space using MDS to situate compression within representational geometry (Methods; Fig. 2d). This embedding enabled a vector decomposition of the representational space into two components, regularity vectors (control→regular for matched elements) and stimulus vectors (tool→animal within a sequence type). Statistical inference was performed against a participant-wise shuffled baseline, generated by permuting condition labels in the RDMs to preserve the set of between-condition distances while disrupting the mapping between condition pairs and their corresponding vectors (Methods).

We first tested whether regularity vectors were aligned across elements by comparing the angle between the tool and animal regularity vectors with the shuffled baseline (smaller angles indicate stronger alignment). Regularity vectors showed reliable alignment across Exps. 1–3 (observed: 66.28*^◦^*± 51.85*^◦^*, 62.37*^◦^* ± 44.59*^◦^*, 78.58*^◦^* ± 31.30*^◦^*; shuffled: 89.83*^◦^* ± 1.80*^◦^*, 89.75*^◦^* ± 1.53*^◦^*, 89.56*^◦^* ± 2.17*^◦^*; paired *t* tests, *t*s ≥ 2.44, *P*s ≤ 0.019, |*d_z_*|s ≥ 0.33; Fig. 2e), indicating a shared regularity axis across distinct elements. In parallel, stimulus vectors were aligned across sequence types (observed vs. shuffled angles in Exps. 1–3: 38.68*^◦^*± 39.60*^◦^* vs. 90.22*^◦^* ± 2.00*^◦^*, 35.57*^◦^*± 30.74*^◦^* vs. 90.00*^◦^*±1.63*^◦^*, and 30.86*^◦^*±24.13*^◦^* vs. 90.26*^◦^*±1.47*^◦^*; paired *t*s ≥ 9.83, *P*s *<* 0.001, |*d_z_*|s ≥ 1.31; Fig. 2f), consistent with preserved category structure. Finally, the regularity axis was more separated from the stimulus axis than expected by chance, quantified as the mean of the four interior angles between regularity and stimulus vectors (observed: 78.89*^◦^* ±17.01*^◦^*, 82.37*^◦^* ±15.46*^◦^*, 84.18*^◦^* ±12.38*^◦^*; shuffled: 60.01*^◦^* ±0.73*^◦^*, 60.06*^◦^* ±0.67*^◦^*, 60.01*^◦^* ± 0.83*^◦^*; paired *t*s ≥ 8.30, *P*s *<* 0.001, *d_z_*s ≥ 1.10; Fig. 2g). Together, these geometric results indicate that sequence-regularity information aligns across elements while remaining distinct from stimulus-related organization in representational space.

Regular-versus-control comparisons of stimulus-vector lengths provided convergent geometric evidence for representational compression of sequence regularity. Tool–animal distances were shorter in regular than in control sequences, reflected by positive control-minus-regular distance differences that exceeded the shuffled baseline (observed Z-scores: 1.48 ± 1.30, 1.83 ± 1.00, 2.97 ± 0.65; shuffled: 0.00 ± 0.05, 0.00 ± 0.06, 0.00 ± 0.07; paired *t*s ≥ 8.51, *P*s ≤ 0.001, *d_z_*s ≥ 1.13; Fig. 2h). This compactness is consistent with compression, indicating that distinct elements within the learned regular sequence occupy a more overlapping region of representational space than their control counterparts.

### Pre-transition compression predicts awareness

We next examined whether representational compression tracks awareness emergence. CCGP indexes cross-element transfer in absolute terms, but its magnitude is constrained by the amount of locally decodable sequence information within each element context. Consequently, identical CCGP values can correspond to different generalization proportions when learners differ in within-element decodability.

We therefore complemented CCGP with a compression rate that normalizes cross-element decoding by matched within-element decoding after centring both on chance (Methods; Supplementary Fig. 4). Operationally, compression rate quantifies the shared above-chance component between within- and cross-element evidence, normalized by within-element decodability with a chance-referenced floor (Methods). This index estimates the proportion of locally decodable sequence information that generalizes across elements. Compression-rate time courses were computed at each time point, and group differences were assessed in non-overlapping 50-ms windows for robust temporal localization.

Time-resolved compression rate showed that transition learners exhibited stronger compression than non-transition learners even before awareness emerged (Fig. 3a; Methods). This pre-transition advantage was confined to late post-stimulus intervals across experiments (0.45–0.50 s in Exp. 1; 0.50–0.60 s in Exp. 2; 0.30–0.40 s and 0.75–0.80 s in Exp. 3). After transition, group differences became broader and more sustained, spanning multiple post-stimulus intervals in each experiment (Exp. 1: 0.10– 0.30 s, 0.45–0.70 s, 0.80–1.05 s, 1.10–1.15 s; Exp. 2: 0.15–0.20 s, 0.25–0.95 s; Exp. 3: −0.10– − 0.05 s, 0.20–0.35 s, 0.45–0.60 s, 1.05–1.20 s). Collectively, these results indicate that a temporally localised compression advantage is already present before transition and is amplified after awareness emerges.

**Fig. 3:**
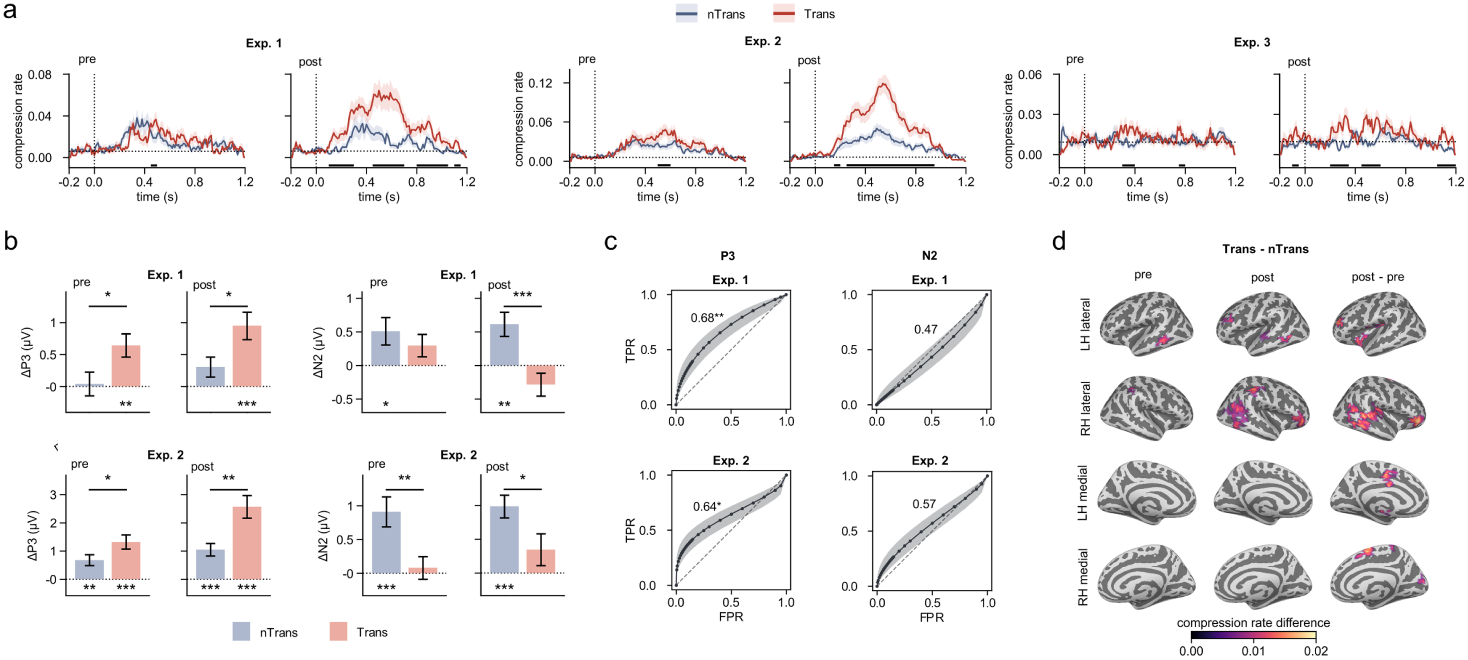
Pre-transition compression predicts subsequent awareness emergence. **a**, Time-resolved compression-rate trajectories in Exps. 1–3, shown separately before and after transition. Lines denote group means for transition learners and non-transition learners, Time 0 marks stimulus onset; shaded bands denote s.e.m., and horizontal black bars indicate 50-ms windows in which Trans exceeded nTrans. **b**, ERP differentiation associated with sequence learning in the two EEG experiments. ΔP3 and ΔN2 were defined as control minus regular amplitudes in component-specific P3 and N2 windows. Bars show group means and error bars denote s.e.m. Asterisks indicate significance for one-sample tests against zero and between-group comparisons. **c**, Fitted ROC analyses testing whether pre-transition compression within P3 and N2 windows predicted later transition status in Exps. 1 and 2. Dashed diagonals indicate chance performance. Shading denotes pointwise s.e. AUC values are shown in each panel, with asterisks indicating significance. **d**, Source-space searchlight maps in Exp. 3 showing compression-rate differences between Trans and nTrans learners before transition, after transition and for the post-minus-pre contrast. Rows show lateral and medial views of the left and right hemispheres. Coloured vertices indicate regions surviving cluster-size permutation correction; colour denotes the Trans–nTrans compression-rate difference.

To interpret the late-stage (>300 ms) compression-rate differences in a time-resolved framework, we analysed event-related potentials (ERPs) evoked by the sequence stimuli in the two EEG experiments (Exps. 1 and 2). ERPs provide well-established temporal markers of sequence processing and offer a univariate complement to MVPA, reducing concerns about shared analytic assumptions. Across conditions, two canonical components were observed: an N2 at Fz peaking at ∼250 ms and a P3 at Pz peaking at ∼500 ms. These component peaks were used to define 100-ms windows for quantifying mean N2 and P3 amplitudes, respectively. ERP waveforms and scalp topographies for each sequence type are summarized in Extended Data Fig. 5.

ERP differentiation associated with sequence learning was quantified as ΔP3 and ΔN2, both defined as control minus regular, given that control sequences were less frequent and less predictable (Fig. 3b). Before transition, transition learners showed a significant ΔP3 effect (Exp. 1/2: 0.64 ± 1.03/1.32 ± 1.40 *µ*V; one-sample *t* tests against zero, *t*s ≥ 3.54, *P*s ≤ 0.001, *d_z_*s ≥ 0.63). This pre-transition effect exceeded the corresponding ΔP3 in non-transition learners (Exp. 1/2: 0.04 ± 0.91/0.67 ± 1.07 *µ*V; *t*s ≥ 2.02, *P*s ≤ 0.048, *g*s ≥ 0.51), and the group difference persisted after transition (transition, Exp. 1/2: 0.95 ± 1.22/2.57 ± 2.23 *µ*V; non-transition, Exp. 1/2: 0.30 ± 0.77/1.05 ± 1.22 *µ*V; *t*s ≥ 2.41, *P*s ≤ 0.019, *g*s ≥ 0.60). In contrast, N2 showed the opposite dissociation: non-transition learners exhibited reliable ΔN2 effects before and after transition (pre, Exp. 1/2: 0.51 ± 1.00/0.91 ± 1.22 *µ*V; post, Exp. 1/2: 0.61 ± 0.89/0.99± 0.92 *µ*V; one-sample *t*s ≥ 2.50, *P*s ≤ 0.020, *d_z_*s ≥ 0.51), whereas transition learners did not (pre, Exp. 1/2: 0.30 ± 0.94/0.08 ± 0.93 *µ*V; post, Exp. 1/2: −0.28 ± 0.96/0.35 ± 1.31 *µ*V; one-sample |*t*|s ≤ 1.77, *P*s ≥ 0.086, |*d_z_*|s ≤ 0.31). Regarding P3 onset, participant-wise P3 latency did not differ significantly between regular and control sequences in any conditions across the two experiments (paired |*t*|s ≤ 1.58, *P*s ≥ 0.124, |*d_z_*|s ≤ 0.29; Supplementary Fig. 6). Thus, we observed a double dissociation in which regular–control differentiation was expressed primarily as a P3 amplitude effect in transition learners and as an earlier N2 amplitude effect in non-transition learners.

We tested whether pre-transition compression within either P3 or N2 peak windows can predict subsequent awareness transition. For each learner, we extracted the peak pre-transition compression rate within the 100-ms P3 and N2 peak windows and evaluated its ability to classify transition status. Pre-transition compression within the P3 peak window predicted subsequent transition in both EEG experiments (AUC [95% CI] = 0.68 [0.53, 0.82] and 0.64 [0.50, 0.78] in Exps. 1 and 2, respectively; *P*s _one-sided_ ≤ 0.036; Fig. 3c, left), whereas prediction was absent for N2-peak-window compression (AUC = 0.47 [0.31, 0.64] and 0.57 [0.41, 0.72] in Exps. 1 and 2, respectively; *P*s _one-sided_ ≥ 0.168; Fig. 3c, right). These results indicate that compression predictive of later awareness emergence is expressed in a P3-aligned time window.

### Cortical compression and predictive information transfer

To localise cortical correlates of representational compression, we performed source-space searchlight decoding in the MEG experiment (Exp. 3). MEG signals were reconstructed on the cortical surface using an LCMV beamformer, and searchlight decoding was applied to source activity within the significant CCGP time window (0.32–1.06 s). At each searchlight, we computed compression rate from cross- and within-element decoding using the same definition as in sensor-level analyses. Com-pression was assessed within each state (pre- and post-transition) relative to the pre-stimulus baseline window (0.2 s; Methods). We then contrasted transition and non-transition learners within each state and tested the corresponding change in group difference across states (post−pre).

In the pre-transition state, both groups showed significantly greater compression relative to baseline in posterior temporal–parietal cortex (Extended Data Fig. 6a–b), indicating that compressive coding of sequence regularity is already present prior to explicit reportability. Critically, transition learners showed a stronger pre-transition compression advantage than non-transition learners, peaking in left middle temporal cortex (lMTG) and also evident in right inferior parietal cortex (Fig. 3d, left). After transition, compression additionally emerged in left rostral middle frontal gyrus (lRMFG) in transition learners (Extended Data Fig. 6c) but remained absent in non-transition learners (Extended Data Fig. 6d), and the transition–non-transition contrast correspondingly extended into lRMFG while the lMTG advantage persisted (Fig. 3d, middle and right). Regional extents and peak coordinates of the searchlight clusters are summarized in Supplementary Table 4. Together, these results indicate that awareness emergence is preceded by a pre-transition posterior temporoparietal compression signature—peaking in lMTG—that becomes more widely expressed, including in prefrontal cortex, after the transition.

Motivated by the spatial expansion from a focal pre-transition lMTG peak to a broader post-transition compression network, we next asked whether pre-transition directed communication predicts subsequent awareness emergence. We defined an ROI network from the Exp. 3 source-space compression maps in transition learners (Extended Data Fig. 6a,c), using the pre-transition lMTG peak as the seed and non-overlapping post-transition compression clusters as targets (Fig. 4a; Supplementary Table 5), including lRMFG, right IFG pars triangularis (rIFGtri), right pars opercularis (rIFGop), right supramarginal gyrus (rSMG), right inferior parietal cortex (rIP), and bilateral parahippocampal gyrus (lPHG and rPHG).

**Fig. 4:**
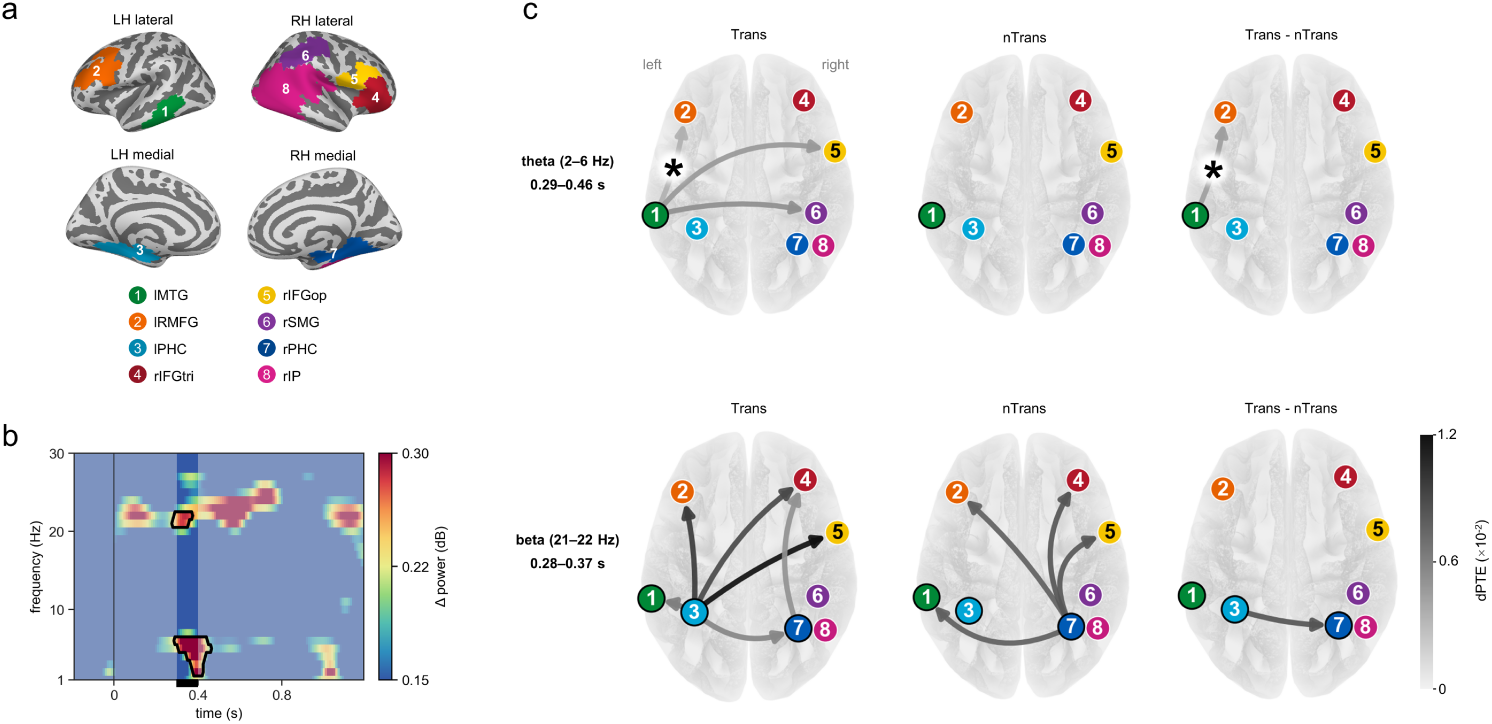
Pre-transition information transfer. **a**, ROI network for dPTE analyses. The pre-transition lMTG compression cluster was used as the seed, and non-overlapping post-transition compression clusters were used as target ROIs. Numbered nodes 1–8 denote lMTG, lRMFG, lPHC, rIFGtri, rIFGop, rSMG, rPHC and rIP, respectively. **b**, Time-frequency power contrast in the lMTG seed, computed as Trans(pre-post) - nTrans(pre-post) to prioritize pre-transition activity in learners who subsequently developed awareness. Black contours mark significant positive clusters that overlapped the 0.30–0.40 s pre-transition Trans *>* nTrans compression window, defining theta (2–6 Hz) and beta (21–22 Hz) bands for subsequent dPTE analyses. Time 0 marks stimulus onset. **c**, Directed phase transfer entropy (dPTE) networks in the theta and beta bands. In the Trans and nTrans columns, arrows mark significant transfer in regular trials; in the Trans–nTrans column, arrows mark significant Trans *>* nTrans differences in regular-trial transfer. Edge colour denotes centred dPTE effect magnitude, with darker edges indicating stronger transfer. Asterisks mark displayed links that additionally showed greater transfer for regular than control trials.

Because directed communication is frequency-specific, we constrained the pre-transition transfer bands in a data-driven manner. Time–frequency power in the lMTG seed, with pre-transition regular and control trials pooled, revealed two significant clusters overlapping the pre-transition compression-rate window (0.30–0.40 s): theta (2–6 Hz) and beta (21–22 Hz; Fig. 4b). We then quantified directed phase transfer entropy (dPTE, centred to [−0.5, 0.5] with 0 indicating no directional bias) for each ordered ROI pair using the same pre-transition trial set as in the compression analyses (Methods), denoting direction as dPTE*_A→B_*. We defined effective transfer as positive dPTE_regular_ that exceeded dPTE_control_, and tested whether this regular–control effect was larger in transition than non-transition learners. This procedure targeted pre-transition directed transfer that was both rule-specific and predictive of the subsequent emergence of awareness.

Theta-band dPTE identified a directed transfer from the lMTG to the lRMFG that satisfied both the rule-specific and transition-predictive criteria (Fig. 4c, top). We first examined positive dPTE in regular trials. In transition learners, the lMTG showed positive dPTE to lRMFG, rIFGop, and rSMG (*t*s ≥ 1.92, *P*s _one-sided_ ≤ 0.033, *d_z_*s ≥ 0.37); these transfers were not significantly above zero in non-transition learners (*t*s ≤ 1.37, *P*s _one-sided_ ≥ 0.091, *d_z_*s ≤ 0.27). We then applied the rule-specificity criterion. Among these candidate pathways, only dPTE_lMTG*→*lRMFG_ was greater in regular trials (3.4 × 10*^−^*^3^) than in control trials (dPTE = −1.3 × 10*^−^*^3^; *t*_26_=1.82, *P*_one-sided_=0.040, *d_z_* = 0.35). This regular–control effect was also larger in transition than non-transition learners (dPTE_mean difference_ = 6.9 × 10*^−^*^3^, Welch *t*_50.0_=1.90, *P*_one-sided_=0.031, *g* = 0.52).

In contrast, beta-band dPTE from the lMTG seed showed no effective directed transfer (*t*s ≤ 0.93, *P*s _one-sided_ ≥ 0.181, *d_z_*s ≤ 0.18; Fig. 4c, bottom). To avoid over-interpreting a null effect tied to a single seed location, we broadened the seed selection to the other two temporal ROIs, lPHG and rPHG. This analysis revealed widespread beta-band transfer in regular trials. In transition learners, positive beta-band dPTE was observed from the lPHG to rPHG, lMTG, lRMFG, rIFGtri, and rIFGop, with an additional rPHG-to-rIFGtri pathway (*t*s ≥ 1.87, *P*s _one-sided_ ≤ 0.036, *d_z_*s ≥ 0.36). In non-transition learners, positive beta-band dPTE was observed from the rPHG to lMTG, lRMFG, rIFGtri, and rIFGop (*t*s ≥ 2.18, *P*s _one-sided_ ≤ 0.020, *d_z_*s ≥ 0.44). However, none of these pathways in transition learners satisfied the rule-specificity criterion (*t*s ≤ 1.37, *P*s _one-sided_ ≥ 0.092, *d_z_*s ≤ 0.26), and the regular–control effect did not differ between transition and non-transition learners (Welch *t*s ≤ 1.51, *P*s _one-sided_ ≥ 0.069, *g*s ≤ 0.41). Thus, beta-band transfer reflected broader communication involving medial-temporal regions rather than a regular-sequence-specific or transition-predictive signal of awareness emergence.

## Discussion

Emergence of awareness during implicit sequence learning makes tacit knowledge avail-able for verbal report and social transmission [8, 27]. Yet despite longstanding work on implicit and explicit sequence learning [10, 13, 15, 28–31], a central mechanistic question remains open: what change in neural representation renders a learned regularity consciously accessible? We addressed this question through representational geometry, asking whether sequence knowledge is organized into an abstract code that supports generalization and conscious access. Our evidence suggests that sequence regularity is compressed into an element-general representation before it becomes reportable, and that explicit access depends on embedding this compressed structure within a broader cortical communication architecture. On this view, conscious access to abstract knowledge reflects a representational transition—compression into a shared neural space coupled with network-level integration—rather than a mere strengthening of the same latent trace.

Neural representations of abstract rules have been documented in language learning [32], reasoning [16], and social learning [33]. In sequence learning, behavioural work has shown robust generalization across exemplars [19, 22, 34–37], but the corresponding neural format has remained difficult to specify. Across Exps. 1–3, cross-element CCGP isolated such a format. Classifiers trained to distinguish regular from control contexts under strict matching of perceptual category and ordinal position generalized to other triplet elements, revealing an element-general code for regularity. Geometry analyses further showed that this code was compressive without collapsing perceptual structure (Fig. 2d–g). RDMs revealed reduced between-category dissimilarity within regular sequences relative to controls (Extended Data Fig. 4), whereas MDS decomposed the embedding into separable axes: a regularity axis (control→regular) aligned across elements beyond shuffled baselines, and a stimulus axis (tool→animal) preserved across sequence types. Critically, these axes were more dissociated than expected under label shuffles (Fig. 2d–h). Thus, compression reflects a geometric transformation that concentrates rule information into a shared representational direction while preserving decodable category structure. This extends classic sequence-learning signatures [38] beyond item-specific associations and echoes compression-like geometries observed in artificial systems [39].

A central message of our results is that representational compression is already present while sequence knowledge remains implicit. Classic accounts of implicit sequence learning have emphasized relatively inflexible perceptual–motor procedural associations [40, 41]. More recent behavioural evidence suggests that implicitly acquired structure can transfer beyond trained instances [42], consistent with abstraction over rule-like regularities [19]. Our data provide direct neural support for this view. In the pre-transition state, CCGP revealed that regularity information generalized across triplet elements differing in perceptual category and ordinal position (Extended Data Fig. 3b), demonstrating an element-general code that cannot be reduced to stimulus identity, position, or motor output. This compressive format was detectable before transition and was present in both transition and non-transition learners across all three experiments, despite differences in task demands, including report versus no-report designs and motor-response versus observation-only learning. Thus, unconscious sequence knowledge is not confined to item-specific sensorimotor bindings; latent structure can already be organized into a compressed, element-general neural format while remaining outside conscious report.

If compression can precede awareness, the critical question becomes which aspect of compression is tied to conscious access. Prior work has often quantified abstraction using CCGP [16, 21, 23, 24, 43–45], but CCGP alone conflates cross-element trans-fer with the strength of sequence information locally decodable within each element context. Similar CCGP values can therefore reflect efficient generalization of weak local evidence or inefficient generalization of strong local evidence. To dissociate these regimes, we introduced a compression rate that normalizes cross-element decoding by matched within-element decoding, providing a time-resolved measure of generalization efficiency. This metric indexes how efficiently locally available sequence evidence is expressed in an element-general code, and is therefore more informative about the representational transformation that supports conscious access than representational strength alone.

Viewed through this efficiency lens, compression predicted conscious access with marked temporal specificity. Transition learners showed a selective pre-transition enhancement of compression in a late post-stimulus window (Fig. 3a). Element-evoked responses in EEG-based Exps. 1–2 further revealed a component-level dissociation in 100-ms peak-window mean amplitudes: in the pre-transition state, transition learners expressed the regular–control effect predominantly in the 100-ms P3 peak window, whereas non-transition learners expressed it earlier in the 100-ms N2 peak window (Fig. 3b; Extended Data Fig. 5). Critically, only P3-window pre-transition compression predicted subsequent transition; N2-window compression did not (Fig. 3c). This P3-window effect was not explained by P3 latency, which showed no regular–control difference across conditions or experiments (Supplementary Fig. 6). This convergence suggests that compression becomes functionally relevant for conscious access during a P3-stage process [46–48]. Beyond its established links to context updating and working-memory operations [49, 50], P3 modulation may therefore index, at least in SRT tasks, the mapping of distinct events onto a shared representational format that supports conscious access.

Exp. 3 extended this account to cortical sources and network dynamics using MEG source imaging. Theoretical accounts have emphasized the close relationship between information flow and the organization of neural representations [51]. Consistent with this view, higher pre-transition compression efficiency in transition learners localised to the left middle temporal gyrus (lMTG). After transition, compression involved a broader frontotemporal network, including lMTG and PFC, consistent with the role of PFC in flexible conscious access and reportable structural knowledge [52]. Using only pre-transition data, directed connectivity analyses further showed theta-band dPTE from the left temporal compression region to PFC targets in the pre-transition state. This finding is consistent with the idea that theta rhythms provide a long-range coordination channel through which information can become available for consciousness-relevant processing and context updating [8, 9, 53, 54]. By con-trast, beta-band interactions formed a broader network, especially involving bilateral medial temporal regions in regular trials, but were not specifically tied to regularity awareness. This dissociation accords with evidence that medial temporal activity during sequence learning can support sequence representations without necessarily tracking conscious awareness [55, 56], and with proposals that beta-band communication supports local representational maintenance or task-state stabilization rather than a regularity-specific route to awareness [8, 57–60].

These source and connectivity results converge with previous SRT evidence showing pre-transition theta dPTE during the emergence of explicit knowledge, together with beta-band activity that was insensitive to sequence regularity [8]. They also differ from classic spatial SRT studies that emphasize parietal sources [37, 61, 62]. This difference is theoretically informative. Spatial SRT regularities are naturally represented in parietal regions that encode spatial position, whereas Exp. 3 removed spatial contingencies and motor requirements and embedded regularity at the level of object categories. Under these conditions, compression localised to left temporal cortex, a region implicated in conceptual and category-level representations [63, 64]. Thus, the cortical origin of predictive routing may depend on where the task-relevant regular-ity is represented, whereas theta-mediated long-distance communication may provide a flexible network mechanism linking compressed representations to conscious access [65].

Theoretically, this study contributes to consciousness research by specifying how an initially implicit regularity becomes formatted for report. The implicit-to-explicit transition in sequence learning has been proposed as a useful window onto conscious access [3, 8]. Our results suggest that awareness is associated with element-general compression of learned structure and its integration with frontal systems through directed theta-band feedforward transfer. This account is broadly consistent with global workspace theories, which emphasize that information becomes reportable when it is made available to a large-scale posterior–prefrontal network [25, 66]. At the same time, substantial compression before awareness shows that integrative representations can arise before reportability, a point compatible with information-integration accounts [67, 68]. The dissociation between pre-conscious compressed representations and awareness-specific frontal representations also resonates with higher-order accounts that distinguish first-order representational content from conscious access to that content [69]. Rather than adjudicating among theories of consciousness [70], our findings identify a concrete representational and network transformation through which latent knowledge becomes consciously accessible.

Several additional findings further clarify how abstract sequence knowledge is expressed during learning. Behaviourally, RT provides a useful online marker of when learned structure begins to guide performance (Fig. 1g and h), although it is not itself a direct measure of awareness [8, 9]. This marker is especially useful in no-report designs because it eliminates the need for repeated awareness probes during learning, which may shift learners toward hypothesis testing or incidental learning [71–73]. Neurally, Exp. 3 revealed predictive perceptual coding: after the first or second item in a triplet, the category of the upcoming item was decodable during the blank interval, approximately 600–800 ms before its onset (Supplementary Fig. 5). This pre-stimulus decoding indicates that learned structure can proactively activate the perceptual representation of the next expected event before sensory input arrives. Finally, the effects cannot be reduced to motor preparation or response execution. In Exps. 1 and 2, sequence decoding and compression remained robust after ICA-based removal of typical motor components; in Exp. 3, trial-by-trial motor responses were removed during learning, with only occasional catch trials using randomised category–button mappings. Thus, although perceptual and motor components can both facilitate sequence learning [74], the present effects primarily reflect perceptual and structural representations rather than learned stimulus–response mappings [75]. Overall, our study establishes a link between abstraction and consciousness, suggesting that research on knowledge generalization [76] and consciousness [77] may benefit from closer integration. Understanding representational generalization requires considering the level of consciousness at which it unfolds, and theories of consciousness may need to account for pre-conscious abstraction. Although EEG and MEG are effective for tracking neural representations in healthy humans [78], future work should use intracranial recordings in non-human primates or patients to examine how compression operates at neuronal and deep-brain levels. Causal perturbation of unconscious features could further test their contribution to the emergence of explicit representations.

## Methods

### Participants and grouping

We recruited 94 (61 females; 22.95 ± 3.31 years), 86 (47 females; 21.97 ± 3.04 years), and 79 (46 females; 20.86 ± 2.32 years) right-handed university students for Exps. 1–3, respectively. All had normal or corrected-to-normal vision, no history of neurological or psychiatric disorders, and were naïve to the study. Each participated in only one experiment. The protocol was approved by the Institutional Review Board of Fudan University (Approval No. FDU-SSDPP-IRB-2025-2-140). Participants gave written informed consent and received monetary compensation.

In each experiment, learners were classified as transition or non-transition based on awareness of the trained sequence rule. For state-based cross-element decoding, transition learners were required to contribute at least three sessions in both the pre-and post-transition states, yielding ≥12 usable trials after category- and position-matching for classifier training; otherwise, they were excluded. This excluded 38, 25, and 27 participants in Exps. 1–3, respectively (Extended Data Fig. 1c). Final analytical samples comprised 56 learners in Exp. 1 (transition *N* = 32; non-transition *N* = 24), 61 in Exp. 2 (transition *N* = 31; non-transition *N* = 30), and 52 in Exp. 3 (transition *N* = 27; non-transition *N* = 25).

Sample size was determined a priori using G*Power (v3.1.9.7) [79]. Assuming Cohen’s *d* = 0.78 from prior work [8], 24 participants per group provide 95% power at *α* = 0.05. We therefore targeted sample sizes sufficient to yield at least 24 learners in both transition and non-transition groups.

### Stimuli and procedure

#### Stimuli

Stimuli were greyscale images from three object categories (faces, tools, animals), with three exemplars per category (Fig. 1a); the same stimulus set was used across experiments. The face stimuli were fully synthetic, AI-generated images and did not depict any real individual. No photographs of participants, authors, or other identifiable persons were used as inputs or references. Images were presented on a mid-gray background (RGB = [128, 128, 128]) using PsychoPy (v2023.2.3) [80]. In Exps. 1–2, stimuli were shown on a monitor (1,024 × 768, 144 Hz) at a viewing distance of 0.8–0.85 m; in Exp. 3, stimuli were rear-projected (32-inch screen; 1,920 × 1,080, 144 Hz) at 0.8 m. Stimuli subtended approximately ±3.6◦ visual angle horizontally and vertically.

#### Procedure

##### Sequence learning task

All experiments used a sequence-rule learning task adapted from the serial reaction time (SRT) paradigm [81]. Learners were not informed about the presence of regularities. Each trial comprised a triplet of three sequentially presented greyscale images drawn from face, tool, and animal categories (three exemplars per category; Fig. 1a). Trials began with a fixation cross (0.70–0.85 s), fol-lowed by three images (0.4 s each) separated by variable blank intervals (0.8–1.2 s). Regular and control trials were interleaved in a pseudo-random order (overall regular:control = 3:2; Extended Data Fig. 1a; see Supplementary Methods for scheduling constraints). In Exp. 1, the trained regular sequence was Type I (face–tool–animal). In Exps. 2–3, learners were trained on either Type I or Type II (face–animal–tool), counterbalanced across learners. Control sequences violated the trained regularity while preserving sufficient trials per category and ordinal position for decoding analyses (Fig. 1a; Supplementary Methods and Supplementary Table 1).

Exp. 1 used a three-choice category-response SRT task with eight sessions (50 standard trials per session; 30 regular/20 control). Learners responded to each image with one of three buttons (category–button mapping counterbalanced) and completed a session-wise awareness assessment after each session.

Exp. 2 used the same category-response task but omitted session-wise awareness assessments (no-report learning stream). It comprised nine sessions of 45 trials each (27 regular/18 control; 405 learning trials total). After learning, learners completed a postlearning awareness test.

Exp. 3 minimized motor-learning contributions by adopting an object-viewing procedure. Each of eight sessions comprised 50 standard trials (30 regular/20 con-trol) plus attention-check (catch) trials. In catch trials, participants responded to the most recent stimulus at a randomly selected triplet position (1–3) using a randomised category–button mapping, preventing fixed motor preparation or motor imagery. These trials were excluded from subsequent analyses, and high accuracy (mean = 96.21%) confirmed sustained task engagement. Learners completed a session-wise awareness assessment after each session and the postlearning awareness test after learning.

Before the main task, learners completed practice blocks (including response-mapping practice where applicable) and proceeded only after two consecutive error-free runs. Short breaks were provided after each session and any accompanying awareness report. Full sequence-construction rules, control constraints, and trial-scheduling procedures are reported in Supplementary Methods.

##### Session-wise awareness test

Participants’ awareness of the sequence regularity was assessed using subjective reports in Exps. 1 and 3, administered after each learning session. The assessment followed a three-step procedure (Supplementary Fig. 1). First, participants provided a free, open-ended description of any observations, impressions, or feelings during the learning task. This prompt did not mention the possibility of a hidden rule or sequence structure. Second, if participants spontaneously mentioned regularities or rules, they were asked whether they believed a specific sequence pattern had been present. Third, if they answered affirmatively, they were asked to describe the sequence order in detail. No performance feedback was provided during the assessment. These reports were used for awareness classification as described in the behavioural analyses.

##### Postlearning awareness test

After learning, participants in Exps. 2 and 3 com-pleted a postlearning awareness test [82] to assess structural knowledge of the trained sequence and the conscious basis of that knowledge (Fig. 1d). Triplets were presented with the same sequential event structure as during learning, but no category responses were required during stimulus presentation; instead, a single conformity judgement was made after each triplet. The test included 75 trials, with 25 trials each of the trained regular sequence, the alternative regular sequence and control sequences (Supplementary Table 2). On each trial, participants judged whether a triplet conformed to the trained sequences and then reported the basis of their judgement using one of four attributions: Guess, Intuition, Memory or Rules. Guess indicated a random judgement with no confidence in its correctness, similar to flipping a coin. Intuition indicated some confidence that the triplet did or did not conform to the trained sequences, without knowing the specific basis for that feeling. Memory indicated explicit recognition of whether the sequence had or had not appeared during learning. Rules indicated that participants had discovered and could describe one or more category-order regularities, and used that knowledge as the basis for their conformity judgement. Definitions of the four attributions were displayed on the screen throughout the test.

### Behavioural analyses

#### RT learning effects

RT learning effects were quantified in Exps. 1 and 2, in which learners made category responses during learning. Analyses included responses to all three triplet positions. Responses were considered missing if no response occurred within 1.3 s of stimulus onset; incorrect and missing responses were excluded at the stimulus level. Mean accuracy was 0.97 and 0.94 in Exps. 1 and 2, respectively.

For each learner, mean RTs were computed separately for regular and control trials, and the learning effect was defined as ΔRT = RT_control_ − RT_regular_, with larger positive values indicating faster responses to regular than control trials. For session-wise analyses, ΔRT was computed for each learning session. For state-based analyses, ΔRT was computed separately for pre- and post-transition sessions in transition learners, or for matched pseudo-transition states in non-transition learners (see *Pseudo-transition session assignment*).

#### Transition classification and session detection

Learners were classified as transition or non-transition using experiment-specific awareness criteria. In Exps. 1 and 3, transition learners were those who correctly reported the trained rule in the session-wise verbal assessment, and the transition session was defined as the first session after which the rule was correctly reported. In Exp. 2, transition required rule-based postlearning knowledge (positive rule-attributed AU_trained_ and ΔAU; see *Postlearning awareness score*), and the transition session was estimated by the best-fitting switchpoint model on the session-wise RT learning effect (Supplementary Table 3). All remaining learners were classified as non-transition.

For state-based analyses, transition learners were required to contribute at least three sessions in both pre- and post-transition states; otherwise, they were excluded (see *Participants and grouping*; Extended Data Fig. 1c). In Exps. 1 and 3, learners transitioning in the final two scheduled sessions completed two additional sessions to ensure sufficient post-transition data.

#### Pseudo-transition session assignment

Non-transition learners served as a baseline for time-on-task effects unrelated to awareness emergence. Because these learners have no observed transition session, we aligned their data to the empirical transition timing of the transition group within each experiment. Concretely, for each transition-session position *T* observed among transition learners, we partitioned *every* non-transition learner’s sessions into “pre-*T*” and “post-*T*” states using the same boundary, computed the corresponding state-based measures, and obtained one non-transition estimate matched to that boundary. Repeating this procedure over all observed *T* values yields a set of matched pre–post estimates for each non-transition learner, which were then averaged within learner and weighted by the empirical frequency of each *T* in the transition group. This yields a single pre/post estimate per non-transition learner aligned to the transition timing distribution. This alignment was applied in all transition–non-transition pre/post comparisons.

#### RT switchpoint modelling

We used Bayesian single-switchpoint modelling to characterize abrupt changes in the RT learning effect (ΔRT = RT_control_ − RT_regular_) across learning sessions in Exps. 1 and 2. For each learner, we fitted a no-switchpoint model and a set of single-switchpoint models in which each session (from the first to the last) was treated as a candidate change point.

Model comparison was based on the widely applicable information criterion (WAIC) [83], where lower WAIC indicates better fit. For each learner, WAIC values were Z-scored across all fitted models (the no-switchpoint model and all candidate switchpoints), so that Z-scored WAIC reflects the relative fit of a model against the alternatives within that learner. RT switchpoint evidence used in Fig. 1g,h was defined as Z-scored WAIC_no-switchpoint_ − mean(Z-scored WAIC_switchpoint_), such that larger values indicate stronger evidence for a switchpoint relative to no switchpoint. Model specification, priors, and sampling procedures are provided in Supplementary Methods.

In Exp. 1, the verbally reported transition session provided an independent reference. The switchpoint model fixed at the report-defined transition session showed superior fit relative to the no-switchpoint model and models placing the switchpoint at other (non-transition) sessions (Extended Data Fig. 2c), providing convergent validation of the model-detected switchpoint. Accordingly, in Exp. 2, the transition session was defined as the session associated with the best-fitting switchpoint model.

#### Postlearning awareness score

Postlearning structural knowledge was quantified using attribution utility (AU), which measures each attribution category’s signed contribution to above-chance discrimination. AU was computed separately for each attribution (*Guess*, *Intuition*, *Memory*, *Rules*) and for each regular-sequence type (trained and alternative).

For each regular-sequence type, its trials were pooled with the same set of control trials to define a two-alternative discrimination. Endorsing the regular sequence (trained or alternative) as conforming to the learned rule and rejecting control sequences as non-conforming were both scored as correct. Treating “yes” responses to the *alternative* regular sequence as correct ensures that AU_alternative_ captures discrimination of rule-like sequences from controls during the test (including test-induced learning or rule generalization) in the same direction as AU_trained_. This design enables a direct comparison between trained and alternative sequences and motivates ΔAU = AU_trained_−AU_alternative_ as a trained-selective structural-knowledge index that controls for test-related contamination.

For each attribution category *i*, we quantified the signed deviation from chance as

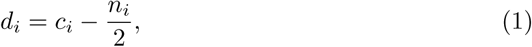

where *c_i_* is the number of correct judgements assigned to attribution *i* and *n_i_* is the total number of trials assigned to that attribution. Attribution utility was then defined as

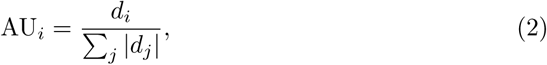

and was set to 0 for all attributions when 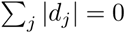

We computed AU*_i_* for the trained and alternative regular sequences and defined ΔAU*_i_* = AU*_i,_*_trained_ − AU*_i,_*_alternative_ as the primary postlearning knowledge scores.

### Neural recording and analyses

#### EEG acquisition and preprocessing

EEG in Exps. 1–2 was recorded with a 128-channel ANT Neuro system (EEGO EE-225; Hengelo, The Netherlands) at 1,000 Hz, using CPz as the online reference and a ground electrode on the nuchal midline. Four EOG electrodes were placed at the outer canthi and above and below the left eye. Electrode impedances were kept below 5 kΩ. Preprocessing was performed in EEGLAB (v2023.1) [84]. Continuous EEG recording was visually inspected to exclude non-stereotyped artefactual segments containing transient muscle activity, jaw-clenching artefacts, or movement-related noise. The retained continuous data were band-pass filtered at 1–80 Hz and notch filtered at 50 Hz. Noisy EEG channels were identified using spectral-based rejection in `pop_rejchan` (1–80 Hz; z-threshold = 2) and removed (Exp. 1: 1.05% ± 1.42% of channels; Exp. 2: 0.65% ± 1.05%). Data were then re-referenced to the common average and down-sampled to 200 Hz.

ICA was then computed with Picard [85] after PCA-based dimensionality reduction to the data rank (retaining 99% variance). To facilitate component classification, IC activations were epoched time-locked to stimulus onset and submitted to ICLabel [86]; labels were then verified by visual inspection of IC time courses, spectra, and scalp maps, including stimulus-locked ERP/topographies, before removal. On average, 2.79 ± 0.97 components were removed in Exp. 1 and 4.43 ± 1.85 in Exp. 2, targeting ocular, cardiac, and head-movement artefacts. Response-related motor components were additionally identified and removed as described in *EEG motor-component removal*.

After artefact and motor-component removal, data were epoched time-locked to stimulus onset from −0.2 to 1.2 s with a −0.2 to 0 s baseline. Epochs with peak-to-peak amplitudes exceeding ±100 *µ*V at any channel were rejected (Exp. 1: 1.63% ± 3.97% of epochs; Exp. 2: 2.14% ± 3.64%). Epochs associated with incorrect button responses were also excluded from decoding (Exp. 1: 4.64% ± 2.24%; Exp. 2: 4.04% ± 2.52%). Previously removed channels were then interpolated using spherical splines [87] on the retained epochs for decoding and ERP analyses.

#### EEG motor-component removal

In Exps. 1 and 2, which required right-hand category responses, motor-related components were removed to minimize motor contamination. We ran an additional full-rank ICA on the cleaned continuous data (no PCA reduction). Candidate motor ICs were identified from response-locked IC activations by aligning to button-press onset (time 0) and inspecting their ERP time courses and scalp topographies. Components were removed if they showed contralateral (left fronto-central/sensorimotor) dominance consistent with right-hand responses (stronger deflection over C3 than C4) and a response-locked extremum with trial-to-trial latency variability *<* 50 ms; the extremum could occur before or after the button press. Selections were visually con-firmed before removal. This procedure removed 1.18 ± 0.39 motor components per learner in Exp. 1 and 1.39 ± 0.67 in Exp. 2. Grand-average ERPs of the removed motor components are shown in Extended Data Fig. 3a.

#### MEG acquisition and preprocessing

Neuromagnetic signals in Exp. 3 were acquired in a magnetically shielded room using a whole-head Elekta Neuromag system (204 planar gradiometers; 102 magnetometers) sampled at 1,000 Hz. EOG and ECG were recorded concurrently for artefact identification. Head position was monitored using four head-position indicator (HPI) coils at the beginning of each session.

MEG preprocessing was performed in MNE-Python (v1.8.0) [88] separately for each session. Bad channels were marked based on Maxwell-based detection (mean ± s.d.: 2.14% ± 0.32% per session). Environmental noise suppression and head-position alignment were performed using temporal signal space separation (tSSS) [89], trans-forming each session to the head position of the first session. Continuous MEG recordings were visually inspected, and non-stereotyped artefactual segments containing transient muscle activity, jaw-clenching artefacts, or movement-related noise were excluded. Data were band-pass filtered at 0.1–150 Hz, notch-filtered at 50/100 Hz, and down-sampled to 400 Hz.

Artefactual components were then removed with FastICA [90] fitted on MEG channels after applying a 1 Hz high-pass filter to improve ICA stability. Candidate components were identified from correlations with EOG signals, aided by ICLabel [86], and confirmed by visual inspection before removal (3.24 ± 1.22 components per session). After preprocessing, all sessions were concatenated; to ensure a consistent channel set, channels marked bad in any session were removed using the union of bad-channel lists across sessions. Data were then epoched relative to stimulus onset (−0.2 to 1.2 s) and baseline-corrected using the pre-stimulus interval (−0.2 to 0 s). Extreme epochs were rejected using peak-to-peak thresholds of 4,000 fT/cm for gradiometers and 5,000 fT for magnetometers (mean ± s.d.: 2.79% ± 8.37%).

#### MEG source reconstruction

MEG source reconstruction was performed in MNE-Python (v1.8.0) [88] using a vector LCMV beamformer [91]. After completing the MEG experiment, each participant underwent a T1-weighted structural MRI scan on a Siemens Prisma 3T scanner (64-channel head/neck coil; Center for MRI Research, Peking University) using an MPRAGE sequence; geometric distortion was corrected using Siemens Syngo (full acquisition parameters are reported in Supplementary Methods).

Individual cortical surfaces were reconstructed with FreeSurfer [92], and an ico4 surface source space was generated for each hemisphere (∼2,562 vertices per hemi-sphere). MEG–MRI coregistration used anatomical fiducials and digitized head-shape points (mean coregistration error: 1.99 ± 0.24 mm per participant). Forward models were computed using a single-layer BEM model (conductivity 0.3 S/m) based on the FreeSurfer-derived inner-skull surface.

Data covariance was estimated from the 0–1.2 s epoch window without shrinkage. Noise covariance was estimated from a same-day 5-min empty-room recording that was preprocessed with the same tSSS, sampling rate, and filtering pipeline as the task data; channels marked bad in the empty-room recording were also removed from the task data to ensure a consistent channel set for covariance estimation. Source time courses were reconstructed with a vector LCMV beamformer using 5% regularization and no weight normalization, and applied to single-trial epochs (−0.2 to 1.2 s). Individual source estimates were then morphed to the fsaverage ico4 space for group-level analyses.

#### Cross-element decoding

Cross-element decoding was used to quantify cross-condition generalization performance (CCGP) [21], measuring the extent to which a decision boundary that discriminates regular from control trials in one element context generalizes to other element contexts. Training was restricted to Positions 2–3, where regular and control trials could be strictly matched for both stimulus category and ordinal position, ensuring that the learned decision boundary captured sequence-structure information rather than perceptual or positional differences. Position 1 was not used for training because sequence structure is not yet expressed until at least two items have been presented. Trained classifiers were then evaluated on held-out epochs from the remaining position(s), including Position 1, to test whether this sequence-structure boundary generalized to contexts not seen during training, consistent with the CCGP framework (Fig. 2a).

Decoding was performed separately for each learner using preprocessed epochs with additional steps optimized for time-resolved classification. Epochs were band-pass filtered at 1–40 Hz and down-sampled to 100 Hz. For each epoch and channel, activity was z-scored relative to the pre-stimulus baseline (−0.2 to 0 s). We used linear logistic regression classifiers (scikit-learn [93]) with L2 regularization, balanced class weights, and the liblinear solver. The inverse regularization strength (*C*) was selected from 0.01, 0.1, 1, 10, and 100 using five-fold cross-validated grid search on the pooled training data [94]. At each time point, features were constructed by concatenating sensor amplitudes from a five-sample temporal neighborhood (two samples before and after the current time point; 50 ms in total), yielding a joint spatiotemporal feature vector for classification [95].

For each learner, two classifiers were trained on the position-specific trained categories (Type I: tool at Position 2 and animal at Position 3; Type II: animal at Position 2 and tool at Position 3). For each time point separately, predicted probabilities and true labels were concatenated across the two train-context configurations, and a single cross-generalization AUC was computed from the pooled predictions. This time-point-wise procedure yielded the CCGP time course.

We applied the same CCGP pipeline across all cross-element decoding analyses, including group comparisons (transition vs non-transition) and both pooled and state-partitioned decoding (transition: pre vs post; non-transition: pseudo-pre vs pseudo-post). Procedures were identical for Exps. 1–2 EEG (all scalp electrodes) and Exp. 3 MEG (magnetometers and gradiometers).

#### Representational dissimilarity matrix (RDM)

Representational dissimilarity matrices (RDMs) [96] provided a decoding-independent characterization of pairwise distances in neural feature space. For direct comparability with CCGP, RDMs were computed from the same preprocessed Position 2–3 epochs and restricted to the experiment-specific CCGP significant time window (Exp. 1: 0.25–1.01 s; Exp. 2: 0.15–1.06 s; Exp. 3: 0.32–1.06 s). Baseline-z-scored activity was averaged across epochs to obtain four condition-specific spatiotemporal patterns (regular–tool, regular–animal, control–tool, control–animal). For each condition, the trial-averaged response formed a time-by-sensor matrix, which was vectorized by concatenating time points and sensors into a feature vector of length *n*_time_ × *m*_sensor_.

Pairwise dissimilarities were computed for all six condition pairs using correlation distance, defined as 1−*r*, where *r* is the Pearson correlation between the corresponding feature vectors. These distances populated a 4 × 4 RDM for each participant (diagonal entries omitted; Extended Data Fig. 4). To emphasize relative distances within each participant, the six off-diagonal dissimilarities were z-scored within participant before group-level visualization and statistical analyses.

#### Multidimensional scaling (MDS)

Multidimensional scaling (MDS) [97] was used to embed the RDM geometry into a low-dimensional space and derive geometric summary metrics for the 2 × 2 design (category: tool vs. animal; sequence type: regular vs. control). The same metric MDS procedure was applied across experiments, using a two-dimensional solution in which Euclidean distances among the four condition points approximated the RDM dissimilarities (z-scored correlation distance, 1 − *r*). MDS was conducted at both the participant and group levels. For participant-level analyses, MDS was fit separately for each participant using the six off-diagonal dissimilarities (z-scored within participant) among the six condition combinations, yielding two-dimensional coordinates for the four conditions. From these coordinates, we defined two vector families that correspond to the CCGP interpretation in the Results: *regularity vectors* (control→regular for tool; control→regular for animal) and *stimulus vectors* (tool→animal within regular; tool→animal within control). We quantified four geometric metrics (reported in Fig. 2e–h): (i) regularity-vector alignment, defined as the angle between the tool and animal regularity vectors; (ii) stimulus-vector alignment, defined as the angle between stimulus vectors in regular and control conditions; (iii) separation between regularity and stimulus axes, quantified as the mean of the four interior angles between regularity and stimulus vectors defined by the quadrilateral formed by the four condition anchors; and (iv) regular-sequence compactness, quantified as the control-minus-regular tool–animal distance, where positive values indicate shorter within-sequence distances for regular than control elements. Statistical inference used participant-wise shuffled baselines. For each participant, condition labels of the z-scored dissimilarities were randomly shuffled 1,000 times, and the full pipeline (MDS embedding and metric computation) was repeated on each shuffled RDM. This procedure preserves the set of dissimilarity values while disrupting their assignment to the six condition pairs, yielding a metric-specific null distribution (e.g., alignment angles within the same vector family approach 90*^◦^* under random labeling). Observed metrics were compared against these shuffled baselines at the participant level for group statistics.

For visualization, MDS was applied to the experiment-level mean RDM obtained by averaging the six z-scored dissimilarities across participants and reconstituting the corresponding group-level RDM (Fig. 2d; Extended Data Fig. 4). Because individual MDS solutions are not uniquely aligned (up to rotation and reflection), group visualization was based on the mean RDM rather than averaging participant-specific coordinate solutions.

#### Compression rate estimation

Compression rate was designed to quantify how efficiently locally decodable sequence information was expressed in a cross-element generalizable format. It jointly considered (i) within-element decoding, which estimated regular–control information available within a fixed element context, and (ii) cross-element decoding, which tested whether this information transferred to a different element context.

Within-element decoding was computed by training and testing regular-versus-control classifiers within the same element context (Position 2 or Position 3), with regular and control trials strictly matched for stimulus category and ordinal position. Training and testing sets were separated by cross-validation. Evidence from the two element contexts was combined to yield a within-element AUC time course (Supplementary Fig. 4). Cross-element decoding was computed as described above for CCGP, using classifiers trained in one element context and tested in another.

For each participant, state, and time point, within- and cross-element AUCs were first expressed relative to chance:

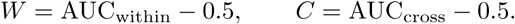

Only above-chance evidence was retained:

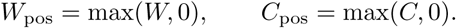

Compression rate quantified the shared above-chance component between local and transferred information, min(*W*_pos_*, C*_pos_), normalized by available local information with a chance floor, *W*_pos_ +0.5 = max(AUC_within_, 0.5). This chance-referenced denominator prevents near-chance within-element decoding from inflating compression estimates. Compression rate was therefore computed as:

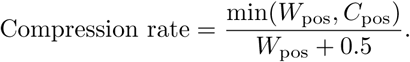

The metric ranges from 0 to 0.5. A value of 0 indicates no above-chance local information or no cross-element transfer, whereas 0.5 corresponds to maximal within- and cross-element decoding (AUC_within_ = AUC_cross_ = 1).

To test whether compression rate within ERP-defined time windows predicted later awareness, we performed ROC analyses using component-specific windows defined from pre-transition ERPs after combining transition and non-transition learners, as described in *Event-related potential analysis*. For each participant, the peak pre-transition compression rate was extracted within the experiment-specific 100-ms P3 or N2 peak window and submitted to separate ROC analyses predicting awareness transition (Fig. 3c).

#### Event-related potential analysis

ERP analyses were conducted in the two EEG experiments (Exps. 1 and 2) to relate the compression-rate effect to canonical ERP components. For ERP analyses only, cleaned epoched EEG data were re-referenced to the average of the left and right mastoids (M1/M2), a conventional reference for visualizing and quantifying midline P3 and N2 components. Stimulus-locked waveforms were then computed at Pz and Fz separately for regular and control trials for each participant (Extended Data Fig. 5). Component windows were defined independently of the regular–control contrast.

For ERP difference analyses, P3 and N2 windows were defined separately for each experiment, group and state after collapsing across regular and control trials. The P3 window was defined as a 100-ms window centred on the Pz peak, and the N2 window was defined analogously from the corresponding Fz waveform. This procedure avoided using the regular–control contrast itself to select the time windows used to test sequence-related ERP effects. For awareness-prediction analyses, pre-transition P3 and N2 windows were defined within each experiment after further collapsing across transition and non-transition learners and across regular and control trials. This yielded one common pre-transition P3 or N2 window per experiment for both groups, avoiding circularity in the transition-versus-non-transition prediction test.

Mean component amplitudes were extracted within the corresponding P3 and N2 windows for regular and control trials. Sequence-related ERP effects were defined separately for each component as ΔP3 = P3_control_ − P3_regular_ and ΔN2 = N2_control_ − N2_regular_ (Fig. 3b). Scalp topographies were averaged within the corresponding P3 and N2 windows (Extended Data Fig. 5).

#### Searchlight decoding

Searchlight decoding was performed in source space in Exp. 3 to map cortical com-pression rate derived from local within- and cross-element decoding. Single-trial vector LCMV source estimates were morphed to fsaverage ico-4 space (5,124 vertices). For each vertex, a local searchlight neighbourhood was defined within the same hemisphere as all vertices within a 10-mm radius. Source activity from neighbouring vertices was concatenated across vertices, the three orthogonal orientation components at each vertex, and time samples within the sensor-level CCGP significant window in Exp. 3 (0.32–1.06 s) to form the classification feature vector.

Within-element and cross-element decoding followed the procedures described in *Cross-element decoding* and *Compression rate estimation*. For source-space searchlight features, PCA retaining 95% of the variance was fitted on the training data within each cross-validation fold and applied to the corresponding test data before logistic regression. Searchlight compression rate was computed at each vertex from the resulting within-element and cross-element AUC values using the formula described in *Compression rate estimation*. State-specific maps were computed separately from trials in the pre-transition and post-transition states for transition and non-transition learners.

Group-level searchlight maps were assessed against the pre-stimulus baseline and between learner groups at each vertex. For each learner, baseline was defined as the mean searchlight compression rate during the −0.2 to 0 s interval. State-specific compression-rate values were tested against this baseline using one-sided one-sample *t*-tests. Transition and non-transition learners were then compared at each vertex using one-sided independent-samples *t*-tests, testing transition *>* non-transition. For post-minus-pre contrasts, state-change maps were first computed within learner and then compared between transition and non-transition learners. For all maps, an initial vertex-wise threshold of *P <* 0.05 was used. Supra-threshold vertices were grouped into spatially contiguous clusters within each hemisphere, and cluster-level correction was performed using 1,000 sign-flip permutations to obtain hemisphere-specific cluster-size thresholds.

#### Time–frequency analysis

Source-space time–frequency analyses were performed in Exp. 3 to identify oscillatory windows for the subsequent directed-connectivity analyses. Because the connectivity analyses tested whether information from the compression source was routed to a broader compression-related network before awareness emerged, ROIs were defined from compression-rate maps in transition learners. The seed ROI was the pre-transition left middle temporal cluster showing significant compression-rate expression. Target ROIs were non-overlapping clusters showing post-transition compression-rate effects in transition learners, including frontal, temporal and parietal regions. The same ROI set was then applied to non-transition learners to provide an anatomical control space (Fig. 4a; Supplementary Table 5).

For each participant and ROI, vector source estimates were reduced to a signed scalar time series using the first principal component of ROI source activity across the full epoch (−0.2 to 1.2 s). Morlet decomposition was applied from 1 to 30 Hz in 1 Hz steps, with *n*_cycles_ = max(0.5 × frequency, 1). Power was computed as the squared magnitude of the complex coefficients, and phase was computed as their angle. Power was multiplied by frequency for 1*/f* scaling and converted to decibels relative to the −0.2 to 0 s baseline.

Frequency bands for phase-transfer entropy analyses were defined from time–frequency power in the left middle temporal seed ROI. To isolate oscillatory activity associated with awareness-related compression rather than general task responses, we contrasted the pre-to-post power change in transition learners with the corresponding pseudo-pre-to-pseudo-post change in non-transition learners, computed as Trans_pre*−*post_ − nTrans_pre*−*post_. Regular and control trials were pooled for this frequency-selection step to avoid biasing the subsequent PTE analysis toward the regular–control contrast. Significant positive time–frequency clusters overlapping the pre-transition Trans *>* nTrans compression-rate window were retained and carried forward to the PTE analyses.

#### Phase transfer entropy

Phase transfer entropy (PTE) was computed from Morlet phase values within the time–frequency clusters identified in the pre-transition time–frequency analysis, using the same trial set as the compression and time–frequency analyses. Transition learners contributed pre-transition trials, and non-transition learners contributed pseudo-pre-transition trials defined from the empirical transition-position distribution. Phase values were discretized into 24 equally spaced bins over [−*π, π*]. For each cluster, the lag was set to one quarter of the oscillatory period at the cluster’s mean frequency.

For each ROI pair, PTE estimated whether the past phase of a candidate source improved prediction of the current phase of a target beyond the target’s own past phase. For the theta cluster, the candidate source was the left middle temporal seed and the targets were prefrontal and parietal ROIs. For the beta cluster, temporal ROIs were additionally tested as candidate sources, as described in the Results.

PTE was computed in both directions for each ROI pair and normalized to obtain centred directional PTE (dPTE), defined as the proportion of source-to-target PTE relative to the summed bidirectional PTE, minus 0.5. Thus, dPTE ranged from −0.5 to 0.5, with positive values indicating preferential source-to-target transfer and 0 indicating no directional bias [98, 99].

For each frequency band and ROI pair, we tested three prespecified effects. First, we tested whether dPTE in regular trials was positive, indicating source-to-target transfer during regular-sequence processing. Second, we tested sequence specificity by comparing dPTE between regular and control trials. Third, we tested transition predictiveness by comparing the regular-minus-control dPTE effect between transition and non-transition learners. One-tailed *t*-tests were used for directional within-group hypotheses, and Welch’s *t*-tests were used for between-group comparisons. Given the prespecified nature of these directional tests, dPTE *P* values are reported uncorrected.

### Statistics of significance

Unless otherwise stated, statistical tests were two-tailed and evaluated at *α* = 0.05. Two-tailed tests were used by default for behavioural comparisons, ERP amplitude analyses, representational-geometry analyses, correlations, group comparisons and all analyses in which deviations in either direction were interpretable. One-sided tests were used only for prespecified directional hypotheses whose expected direction was defined before testing by the analysis metric or by the transition hypothesis, and not by the observed direction of the data. These directional tests included above-chance decoding and ROC effects (AUC *>* 0.5), compression above the pre-stimulus baseline, transition *>* non-transition compression, positive source-to-target dPTE in regular trials, regular *>* control dPTE for sequence-specific transfer, and larger regular-minus-control dPTE effects in transition than non-transition learners. All one-sided tests are explicitly labelled as such in the Results or figure captions.

Participant-level values were used for group-level inference. Data are reported as mean ± s.d. or mean with 95% CI. Error bars denote s.e.m.; ROC shaded bands denote pointwise s.e. For all ROC analyses, AUC confidence intervals and *P* values were estimated using 1,000 bootstrap resamples and 1,000 label permutations, respectively. Time-resolved decoding effects were assessed using cluster-based permutation tests with 1,000 permutations. Temporal clusters in which AUC exceeded chance (0.5) were evaluated with one-sided cluster-corrected tests at *P <* 0.05, because the hypothesis concerned above-chance information rather than any deviation from chance. For state and group comparisons, each learner’s peak AUC or compression-rate value was extracted within the corresponding experiment-specific significant window, unless an ERP-defined window was used.

Behavioural, neural and representational measures were analysed using *t*-tests, mixed ANOVA, chi-square tests, Pearson correlations, ROC analyses, permutation tests or shuffled-baseline tests, as appropriate. Welch’s correction was used for unequal-variance group comparisons, and Bonferroni or Benjamini–Hochberg false-discovery-rate correction was applied where indicated. Representational-geometry analyses were evaluated against participant-wise shuffled baselines. Source-space searchlight maps were corrected within each hemisphere using cluster-size permutation testing with 1,000 random group-label permutations.

Effect sizes are reported where applicable: Cohen’s *d_z_* for one-sample and paired *t*-tests, Hedges’ *g* for independent-samples *t*-tests, partial eta squared 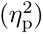 for mixed ANOVA, *ϕ* for 2×2 chi-square tests and Pearson’s *r* for correlations. Permutation, bootstrap and shuffled-baseline procedures used 1,000 iterations unless stated otherwise. Analyses were performed in Python using SciPy, Pingouin, statsmodels, scikit-learn, MNE-Python and custom scripts.

## Acknowledgements

We thank the Meta-Innovation Center at Fudan University for their support in data collection. We thank Zhiliang Yang for his advice and support, and Xiaoqing Hu for discussions. The computations in this research were performed using the Undergraduate Experimental Teaching Supercomputing Plat-form of Fudan University. We thank National Center for Protein Sciences at Peking University in Beijing, China, for assistance with data acquisition, and Baojia Sun, Wenhua Yan, and Weiwei Men for help with MEG and sMRI data collection.

## Declarations

### Data availability

Data used in this study are available at Science Data Bank (https://www.scidb.cn/anonymous/bnl1UTdi).

### Extended data

The Extended Data figures 1–7 are included at the end of the manuscript.

## Extended Data Figures

**Extended Data Fig. 1:**
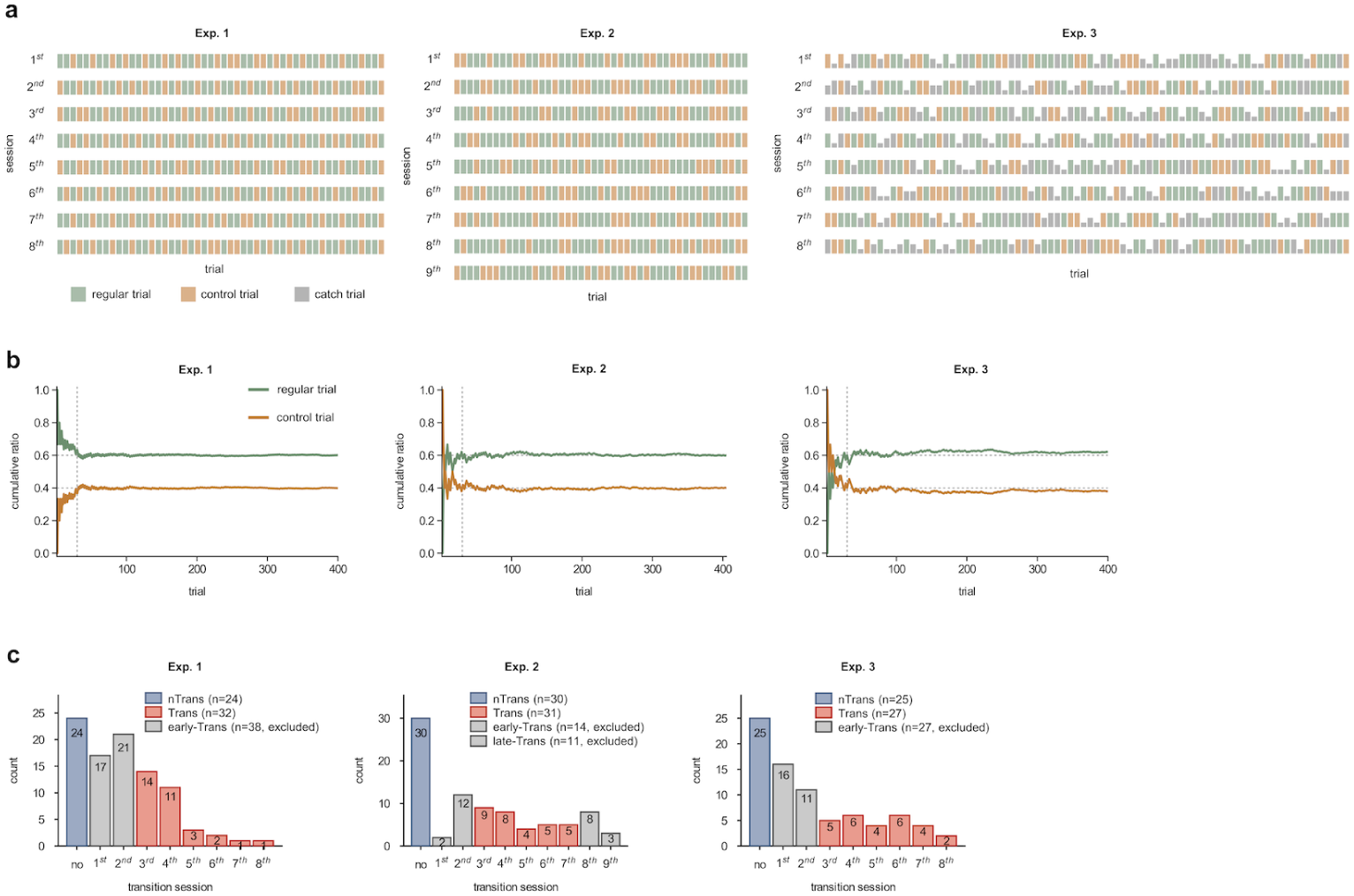
Trial schedule and transition-session distribution. **a**, Trial schedules in Exps. 1–3. Rows depict the chronological order of regular and control trials within each session. In Exp. 3 (right), grey rectangles indicate catch trials; rectangle height denotes the probed item position. **b**, Cumulative proportions of regular and control trials across training. The vertical dotted line marks trial 30, after which the running proportions stabilize near 60% (regular) and 40% (control) (horizontal dotted lines). **c**, Distribution of transition-session positions across Exps. 1–3. Bars show the number of non-transition learners (nTrans), transition learners (Trans), and learners excluded due to early or late transitions that yielded insufficient data for state-based analyses (fewer than three sessions in either the pre- or post-transition state).

**Extended Data Fig. 2:**
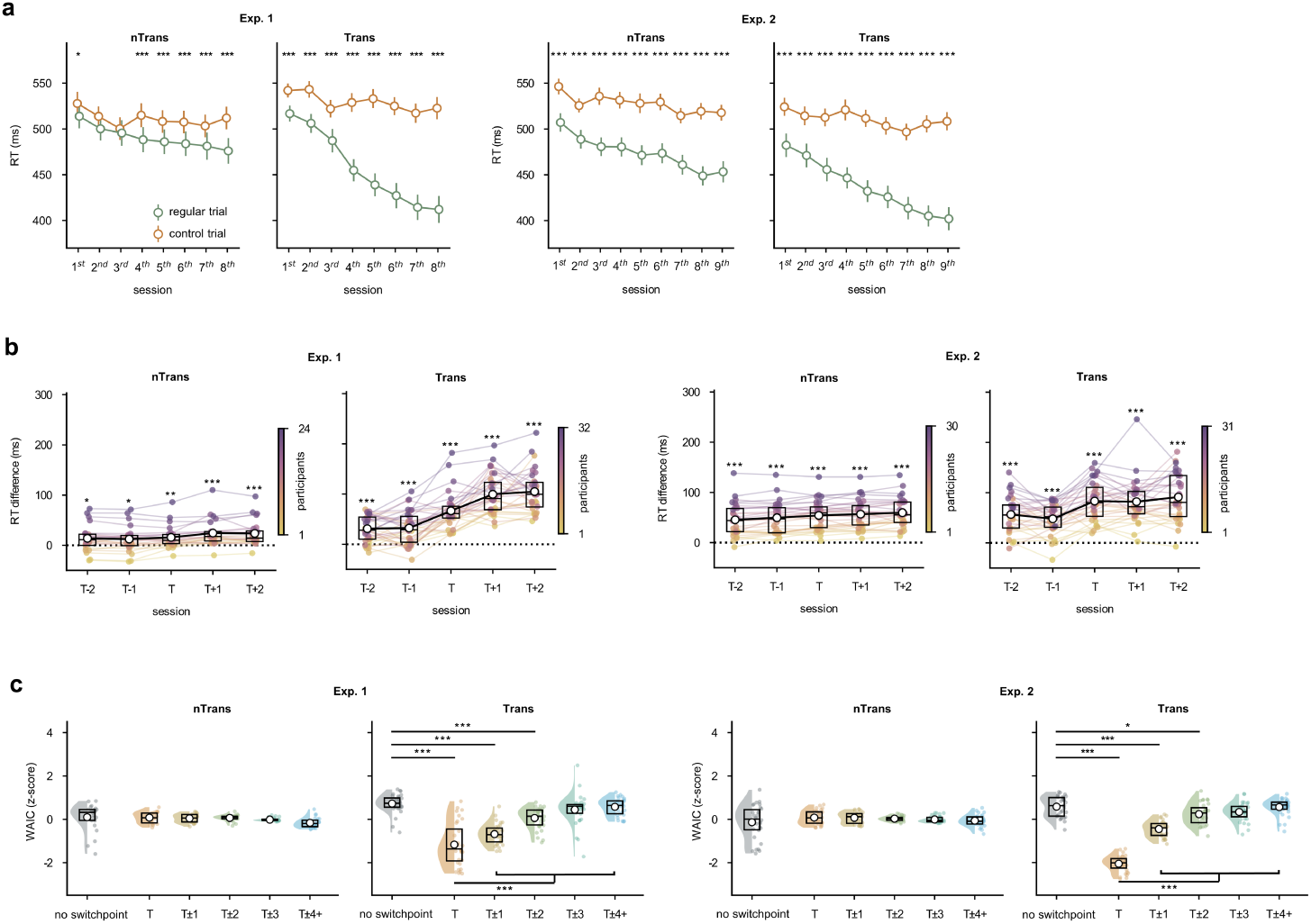
Session-wise RT learning effects and switchpoint modelling. **a**, Session-wise RTs for regular and control trials. Regular RTs were shorter than control RTs in most cases (significant sessions: paired *t*s ≥ 2.54, *P*s ≤ 0.018, *d_z_*s ≥ 0.52), except for non-transition learners in Sessions 2–3 of Exp. 1 (paired *t*s ≤ 2.04, *P*s ≥ 0.053). **b**, Transition-aligned RT learning effect (RT_control_ − RT_regular_). Transition learners were aligned to their observed transition session (T), whereas non-transition learners were aligned to pseudo-transition sessions. The learning effect was reliably above zero across aligned sessions in both groups and experiments (*t*s ≥ 2.32, *P*s ≤ 0.029, *d_z_*s ≥ 0.47). The transition-centred gain, defined as the change from T−1 to T, was larger than all non-T session-to-session gains in transition learners (paired *t*s ≥ 4.38, *P*s *<* 0.001, *d_z_*s ≥ 0.77), but not in non-transition learners (paired *t*s ≤ 0.90, *P*s ≥ 0.374, *d_z_*s ≤ 0.16). **c**, Bayesian switchpoint model comparison for RT learning effects. Z-scored WAIC is shown for a no-switchpoint model and for candidate switch-point models binned by distance from each learner’s transition session (T), or from pseudo-transition sessions in non-transition learners. Lower WAIC values indicate better fit. In transition learners, switchpoint models at or near T outperformed the no-switchpoint model (T, T±1 and T±2: *t*s ≥ 2.27, *P*s ≤ 0.031, *d_z_*s ≥ 0.41), whereas farther models did not (T±3 and T±4+: *P*s ≥ 0.053). In non-transition learners, no candidate switchpoint model differed reliably from the no-switchpoint model (*P*s ≥ 0.057). Among models with switchpoints, the T model outperformed the mean of all non-T models in transition learners (paired *t*s ≤ −6.39, *P*s *<* 0.001, *d_z_*s ≤ −1.13), but not in non-transition learners (paired *t*s ≤ 1.88, *P*s ≥ 0.073, *d_z_*s ≤ 0.38). Coloured lines or points show individual learners and black markers summarize group-level statistics. In box plots, open circles indicate means and horizontal lines indicate medians.

**Extended Data Fig. 3:**
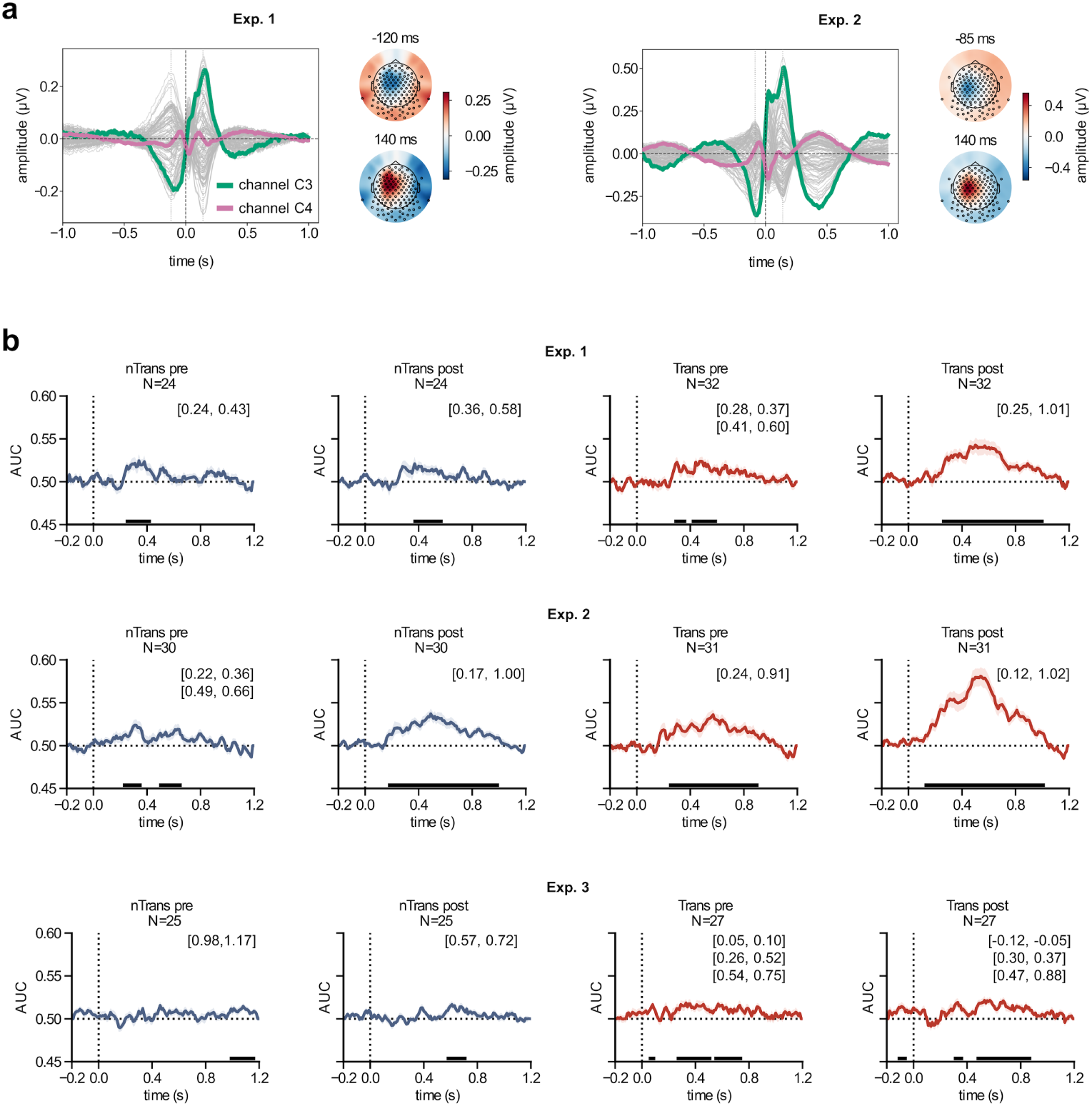
Motor-related ERP characterization and CCGP decoding. **a**, Grand-average response-locked ERPs across all learners and all 128 EEG channels (butterfly plot; time 0 indicates button press). Motor-related components identified from these response-locked signals were removed from the EEG data before decoding analyses. Representative traces at C3 (green) and C4 (magenta) illustrate a lateralized motor potential, with stronger activity over the contralateral (left-central) motor region. Topographies (right) show scalp distributions at the peak global field power before and after the button press, highlighting the expected left-central lateralization. **b**, CCGP time courses for Exps. 1–3, shown separately by group (transition vs non-transition) and state (pre- vs post-transition), corresponding to the analyses in Fig. 2b,c. Time 0 marks stimulus onset; significance was assessed with 1,000 permutations and cluster-based correction (*P <* 0.05); shaded areas denote s.e.m. Significant time clusters are marked along the bottom of each panel and summarized in the panel corner.

**Extended Data Fig. 4:**
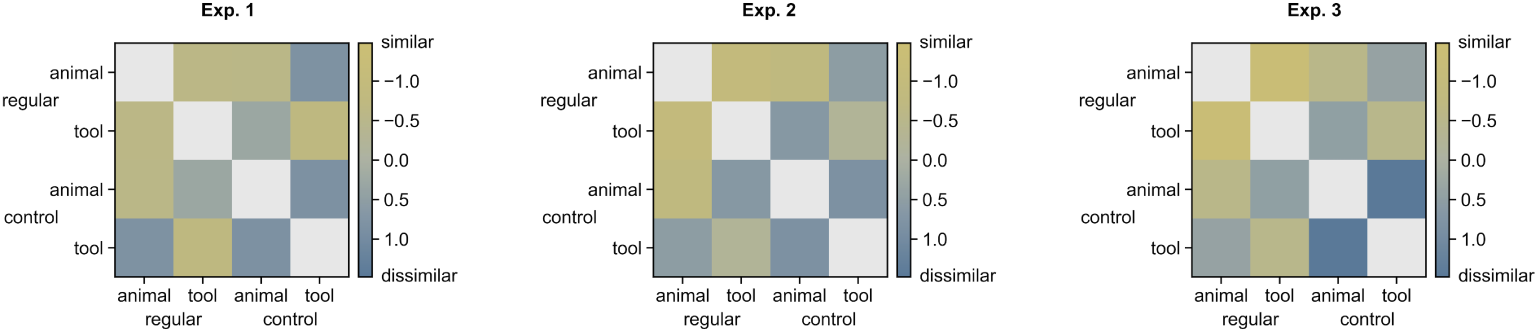
Z-scored representational dissimilarity matrices (RDMs) in Exps 1–3. Neural patterns were extracted from the experiment-specific significant CCGP time window and averaged by condition for each participant. RDM entries were computed as 1 − *r*, where *r* is the Pearson correlation between the sensor-level time series (EEG/MEG) for each pair of the 2×2 conditions (tool vs. animal) × (regular vs. control), yielding a 4×4 dissimilarity matrix. Matrix values were z-scored within participants.

**Extended Data Fig. 5:**
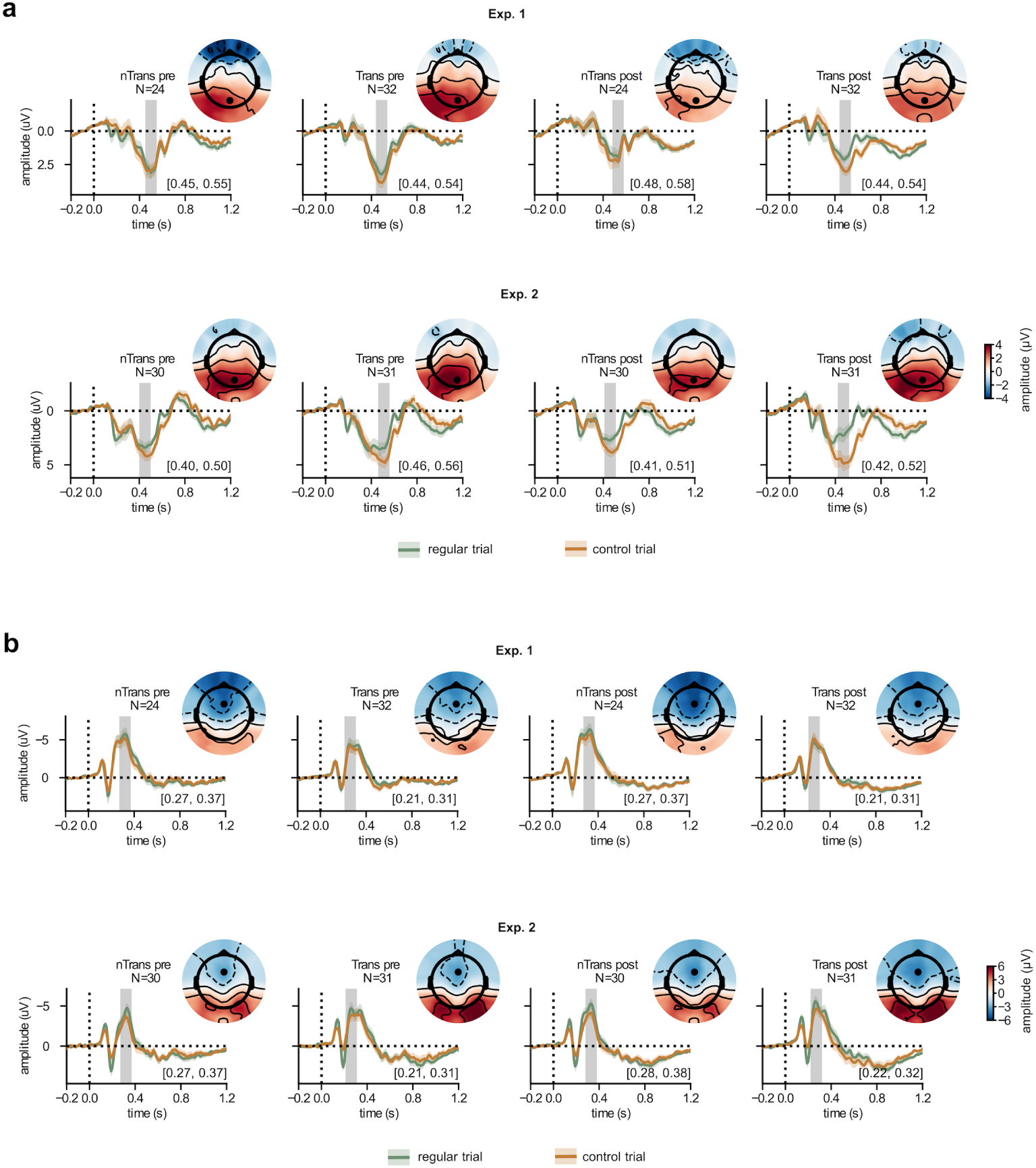
Event-related potentials in the two EEG experiments (Exps 1 and 2). **a**, Pz event-related potential (ERP) waveforms for regular and control trials in Exps. 1 and 2, shown separately for non-transition and transition learners and for pre- and post-transition states. Time 0 marks stimulus onset; shading around each waveform denotes the s.e.m. across participants. Grey shaded windows indicate the independently defined group- and state-specific 100-ms P3 peak intervals used for ERP analyses, with the corresponding time ranges shown in each panel. Scalp topographies show mean P3 amplitudes averaged across regular and control trials within the corresponding P3 peak window and are displayed at the upper right of each condition panel. Channel Pz is marked by a black dot. **b**, Fz ERP waveforms and corresponding N2 scalp topographies, plotted as in **a**. Grey shaded windows indicate the independently defined 100-ms N2 peak intervals used for ERP analyses. Channel Fz is marked by a black dot.

**Extended Data Fig. 6:**
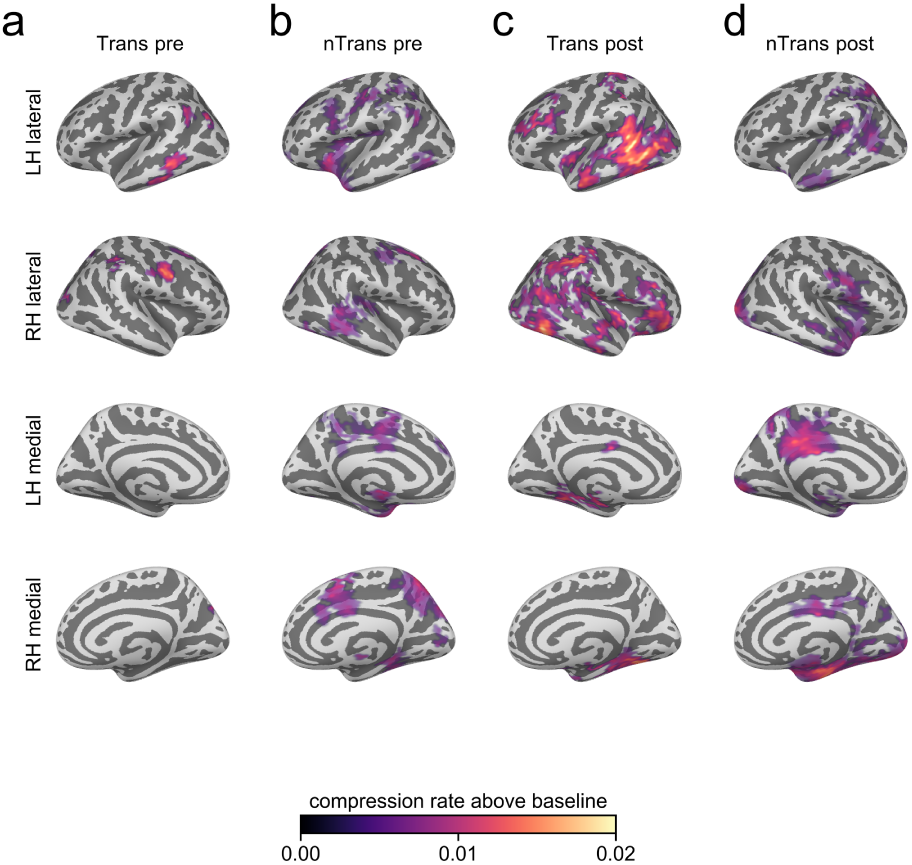
Source-space compression-rate maps above baseline in Exp. 3. Cortical maps show source-space vertices at which the searchlight compression rate exceeded the pre-stimulus baseline in transition and non-transition learners before and after transition (**a–d**). The baseline was defined as the mean compression rate during the MEG pre-stimulus interval (−0.2 to 0 s; baseline value = 5.9 × 10*^−^*^3^). Columns show transition pre-transition (**a**), non-transition pre-transition (**b**), transition post-transition (**c**) and non-transition post-transition (**d**) maps. Rows show lateral and medial views of the left and right hemispheres. At each vertex, compression rate was tested against the baseline value using one-sided one-sample *t*-tests. Supra-threshold vertices (*P <* 0.05) were grouped into connected clusters within each hemisphere, and clusters were retained using hemisphere-specific cluster-size thresholds estimated from 1,000 sign-flip permutations (cluster alpha = 0.10). Colour indicates the mean compression rate above baseline.

**Extended Data Fig. 7:**
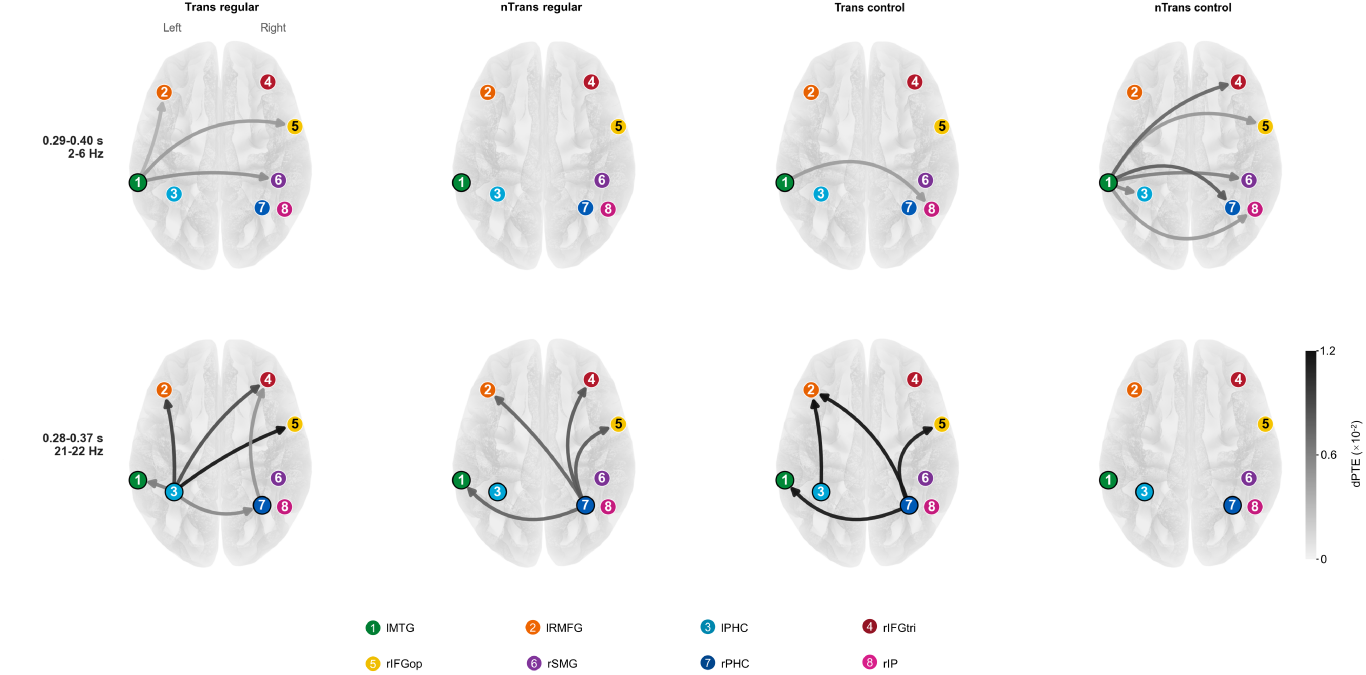
Regular- and control-trial information transfer in the theta and beta windows. Directed phase transfer entropy (dPTE) networks are shown separately for transition learners and non-transition learners, and for regular and control trials. Columns show Trans regular, nTrans regular, Trans control and nTrans control. The top row shows theta-band transfer in the 0.29–0.40 s, 2–6 Hz window, testing outgoing transfer from the lMTG seed (node 1). The bottom row shows beta-band transfer in the 0.28–0.37 s, 21–22 Hz window, testing outgoing transfer from temporal seeds, including lMTG, lPHC and rPHC (nodes 1, 3 and 7). Arrows mark significant positive directed transfer in the corresponding group-by-condition cell, assessed as centred dPTE *>* 0 with one-tailed one-sample tests (*P <* 0.05, uncorrected). Edge colour denotes centred dPTE effect magnitude, with darker edges indicating stronger transfer. Numbered nodes 1–8 denote lMTG, lRMFG, lPHC, rIFGtri, rIFGop, rSMG, rPHC and rIP, respectively.

