## Supplementary Information for "Compressed representations underpin knowledge awareness in sequence learning"

### Supplementary Methods

#### Sequence construction and trial scheduling

Standard learning trials consisted of three sequentially presented stimuli, each drawn from one of three categories: face (F), tool (T) and animal (A). Regular trials followed the trained category-level sequence. In Exp. 1, the trained regular sequence was FTA. In Exps. 2 and 3, the trained regular sequence was either FTA or FAT, counterbalanced across participants. Control trials were constructed to violate the trained regularity while preserving sufficient observations for decoding analyses across stimulus categories and sequence positions. In Exp. 1, control sequences satisfied three constraints (Supplementary Table 1a). First, they did not contain a face in the first position, increasing the distinction between regular and control trials at sequence onset. Second, they did not contain the regular second-to-third sub-sequence, that is, a tool in the second position followed by an animal in the third position. Third, they contained at least one category-position feature shared with the regular sequence, either a tool in the second position or an animal in the third position, to ensure sufficient data for sequence decoding. These constraints yielded eight possible control templates when repeated categories were allowed. Regular and control trials were interleaved pseudo-randomly, with no more than two consecutive trials of the same type. In Exps. 2 and 3, control sequences were defined as category triplets that did not begin with a face (Supplementary Table 1b). Because regular sequences in these experiments were face-leading sequences (FTA or FAT), this rule excluded both the trained regular sequence and the alternative face-leading sequence from the learning phase. The control set therefore contained 18 possible templates, corresponding to two possible first-position categories and three possible categories at each of the second and third positions. Regular and control trials were interleaved pseudo-randomly, with regular-to-control and control-to-regular transition proportions constrained to be approximately 4:6.

#### Switchpoint Bayesian modelling

We used Bayesian model comparison to evaluate whether behavioural learning effects changed abruptly during learning. For each participant, we fitted three classes of models: (1) a no-switchpoint model, which assumed that the learning effects remained stable across learning sessions; (2) a transition-switchpoint model, which placed the switchpoint at the independently defined transition session; and (3) non-transition-switchpoint models, which placed the switchpoint at each candidate session other than T. Comparing these models tested whether a sudden change was present and whether that change was specifically aligned with the transition session.

The switchpoint model was applied to trial-level RTs from Exps. 1 and 2. RTs were log-transformed and linearly detrended within participant before model fitting. For each participant, log-transformed and detrended RTs were modeled separately for control and regular trials. Let $y_{\text{control}}(k)$ and $y_{\text{regular}}(k)$ denote the transformed RT for trial $k$ in control and regular trials, respectively. The model was:

$$y_{\text{control}}(k) \sim \text{Normal}(\mu_{\text{control}}, \sigma^2) \tag{1}$$

$$y_{\text{regular}}(k) \sim \text{Normal}(\mu_{\text{regular}}(k), \sigma^2) \tag{2}$$

$$\sigma \sim \text{HalfNormal}(2S_{\text{obs}}) \tag{3}$$

$$\mu_{\text{control}} \sim \text{Normal}(\bar{x}_{\text{obs}}, 2S_{\text{obs}}) \tag{4}$$

$$\mu_{\text{regular}}(k) = \mu_{\text{control}} - \Delta\mu(k) \tag{5}$$

where $\bar{x}_{\text{obs}}$ and $S_{\text{obs}}$ denote the sample mean and sample s.d. of the participant's log-RTs, respectively. $\Delta\mu(k)$ denotes the learning effect in log-RT units, corresponding to faster responses for regular than control trials when positive.

In the no-switchpoint model, $\Delta\mu(k)$ was constant across all sessions:

$$\Delta\mu(k) = \Delta\mu \tag{6}$$

$$\Delta\mu \sim \text{Normal}(\bar{x}_{\text{obs}}, 2S_{\text{obs}}) \tag{7}$$

In the transition-switchpoint model, $\Delta\mu(k)$ was allowed to differ pre and post the transition session $N$:

$$\Delta\mu(k) = \begin{cases} \Delta\mu_{\text{pre}}, & k \le N \\ \Delta\mu_{\text{post}}, & k > N \end{cases} \tag{8}$$

$$\Delta\mu_{\text{pre}} \sim \text{Normal}(\bar{x}_{\text{obs}}, 2S_{\text{obs}}) \tag{9}$$

$$\Delta\mu_{\text{post}} \sim \text{Normal}(\bar{x}_{\text{obs}}, 2S_{\text{obs}}) \tag{10}$$

For the transition-switchpoint model, $N$ was fixed to the participant's transition session $T$. For non-transition-switchpoint models, the same model was fitted repeatedly with $N$ fixed to each candidate session other than $T$.

Posterior distributions were estimated using Markov chain Monte Carlo with a Metropolis-Hastings sampling scheme. For each model, four chains of 40,000 samples were run. The first half of each chain was discarded as burn-in, and every fifth sample from the remaining samples was retained to reduce autocorrelation. Model fit was compared using widely applicable information criterion (WAIC), with lower WAIC indicating better out-of-sample predictive fit.

#### MRI data acquisition and preprocessing

Structural MRI data were acquired at the Center for MRI Research, Peking University, using a 3-T Siemens Prisma scanner equipped with a 64-channel head-neck coil. Participants lay comfortably on the scanner bed, and their heads were stabilized with foam pads to minimize head motion. High-resolution T1-weighted anatomical images were acquired using a three-dimensional magnetization-prepared rapid gradient-echo

(MPRAGE) sequence with the following parameters: 192 contiguous sagittal slices, repetition time (TR) = 2530 ms, echo time (TE) = 2.98 ms, inversion time (TI) = 1100 ms, flip angle = 7°, receiver bandwidth = 240 Hz per pixel, field of view = 256 mm × 224 mm, matrix size = 256 × 224, slice thickness = 1 mm, and acquisition voxel size = 1 mm × 1 mm × 1 mm. Images were reconstructed with an interpolated voxel size of 0.5 mm × 0.5 mm × 1 mm. The acquisition time was 5 mins 58 seconds. Vendor-provided distortion correction was applied using Siemens syngo software.

Structural images were then further preprocessed before source reconstruction. Bias-field inhomogeneity was corrected using ANTs $N4BiasFieldCorrection$, and image denoising was performed using ANTs $DenoiseImage$. Intensity normalization was performed using FreeSurfer $mri\_normalize$, and preprocessing quality was visually inspected in Freeview. Individual anatomical reconstruction and cortical/head-surface reconstruction were performed with FreeSurfer $recon - all$. The resulting surfaces were used for MEG–MRI coregistration, head-model construction and source reconstruction.

### Supplementary Tables

**Supplementary Table 1: Control sequence assignment for the learning task.**

**a. Control sequences in Exp. 1.**

| | | |
|---|---|---|
| tool–face–animal | tool–tool–face | tool–tool–tool |
| tool–animal–animal | animal–face–animal | animal–tool–face |
| animal–tool–tool | animal–animal–animal | |

**b. Control sequences in Exps. 2 and 3.**

| | | |
|---|---|---|
| tool–face–face | tool–face–tool | tool–face–animal |
| tool–tool–face | tool–tool–tool | tool–tool–animal |
| tool–animal–face | tool–animal–tool | tool–animal–animal |
| animal–face–face | animal–face–tool | animal–face–animal |
| animal–tool–face | animal–tool–tool | animal–tool–animal |
| animal–animal–face | animal–animal–tool | animal–animal–animal |

*Note.* Templates denote category-level sequences, not specific image exemplars. Repeated categories were allowed in control templates. In Exp. 1, control sequences were constrained to exclude faces from the first position, exclude the regular second-to-third sub-sequence, that is, a tool in the second position followed by an animal in the third position, and retain at least one, but not both, category-position feature shared with the regular sequence. This constraint ensured that control trials contained a feature usable for sequence decoding, either a tool in the second position or an animal in the third position, while preventing control sequences from being overly similar to the regular sequence. These constraints yielded eight possible control templates. In Exps. 2 and 3, control sequences were defined as all category-level triplets that did not begin with a face, yielding 18 possible templates.

**Supplementary Table 2**: **Sequence assignment for the postlearning structural knowledge test.**

**a. Regular sequence assignment**

| **Learner type** | **Trained regular sequence** | **Alternative regular sequence** |
|---|---|---|
| Type I trained learners | face–tool–animal | face–animal–tool |
| Type II trained learners | face–animal–tool | face–tool–animal |

**b. Shared control sequences used for both Type I and Type II learners**

| | | |
|---|---|---|
| face–face–face | face–face–tool | face–face–animal |
| face–tool–face | face–tool–tool | face–animal–face |
| face–animal–animal | tool–face–face | tool–face–tool |
| tool–face–animal | tool–tool–face | tool–tool–tool |
| tool–tool–animal | tool–animal–face | tool–animal–tool |
| tool–animal–animal | animal–face–face | animal–face–tool |
| animal–face–animal | animal–tool–face | animal–tool–tool |
| animal–tool–animal | animal–animal–face | animal–animal–tool |
| animal–animal–animal | | |

*Note.* Type I and type II learners were trained on opposite regular sequences. The same 25 control sequences were used for both learner types, corresponding to all possible triplets over face, tool and animal categories ($3 \times 3 \times 3 = 27$) except the two regular sequences.

**Supplementary Table 3: Model-based switchpoint estimates for participants in Exp. 2.**

| Subject | Group | SP1 | SP2 | SP3 | SP4 | SP5 | SP6 | SP7 | SP8 | SP9 | No SP | Best SP |
|---|---|---|---|---|---|---|---|---|---|---|---|---|
| sub01 | Trans | 0.09 | 1.61 | -0.36 | **-2.48** | -0.03 | 0.16 | -0.40 | 0.57 | 0.78 | 0.06 | SP4 |
| sub02 | late-Trans | -0.36 | 0.23 | 0.67 | 0.84 | 1.03 | -0.28 | -0.24 | **-2.51** | 1.00 | -0.38 | SP8 |
| sub03 | Trans | 0.35 | 1.05 | 0.92 | **-2.03** | -0.58 | -1.07 | -0.73 | 1.12 | 0.64 | 0.34 | SP4 |
| sub04 | early-Trans | 0.55 | **-2.73** | -0.74 | 0.02 | 0.30 | 0.10 | 0.65 | 0.80 | 0.49 | 0.55 | SP2 |
| sub05 | late-Trans | 0.54 | 0.24 | 0.86 | 0.72 | 0.67 | 0.27 | -0.49 | -0.72 | **-2.61** | 0.53 | SP9 |
| sub06 | Trans | 1.15 | -0.84 | -0.79 | **-1.71** | -1.12 | 0.18 | 0.43 | 0.31 | 1.24 | 1.15 | SP4 |
| sub07 | early-Trans | 1.14 | **-1.77** | -1.51 | -0.18 | -0.24 | 0.20 | 0.32 | -0.37 | 1.26 | 1.14 | SP2 |
| sub08 | early-Trans | 0.69 | **-2.29** | -1.33 | 0.53 | -0.26 | 0.33 | 0.63 | 1.08 | -0.07 | 0.69 | SP2 |
| sub09 | late-Trans | 0.39 | 0.62 | 0.68 | 0.62 | 0.67 | 0.64 | -0.35 | **-2.46** | -1.20 | 0.39 | SP8 |
| sub10 | Trans | 0.64 | 0.53 | **-1.95** | -1.02 | -1.34 | -0.01 | 0.53 | 0.97 | 1.04 | 0.63 | SP3 |
| sub11 | Trans | 1.08 | -0.10 | -0.51 | -1.27 | **-1.82** | -0.55 | 0.07 | 0.76 | 1.26 | 1.08 | SP5 |
| sub12 | Trans | 0.34 | 0.90 | 0.27 | **-2.76** | -0.60 | 0.81 | 0.09 | 0.12 | 0.51 | 0.32 | SP4 |
| sub13 | early-Trans | -0.13 | **-2.91** | 0.48 | 0.52 | 0.46 | 0.47 | 0.49 | 0.49 | 0.25 | -0.13 | SP2 |
| sub14 | late-Trans | 0.26 | 0.50 | 0.52 | 0.37 | 0.53 | 0.34 | 0.47 | **-2.91** | -0.35 | 0.26 | SP8 |
| sub15 | Trans | 0.87 | 0.13 | 0.19 | **-2.02** | -1.35 | 0.67 | -0.30 | 1.36 | -0.43 | 0.86 | SP4 |
| sub16 | Trans | 0.63 | -1.84 | -0.24 | **-1.86** | -0.02 | 0.29 | 0.49 | 0.78 | 1.17 | 0.62 | SP4 |
| sub17 | early-Trans | 0.15 | **-2.14** | 0.74 | 0.27 | -0.82 | 0.05 | -0.91 | 1.10 | 1.45 | 0.12 | SP2 |
| sub18 | nTrans | 1.36 | 0.48 | 0.61 | 0.29 | 0.06 | -1.09 | -1.41 | **-1.55** | -0.13 | 1.36 | SP8 |
| sub19 | Trans | 1.02 | -0.89 | -0.90 | 0.01 | -0.41 | -1.03 | **-1.36** | 1.27 | 1.27 | 1.02 | SP7 |
| sub20 | nTrans | 0.10 | -0.76 | 0.83 | 1.46 | 0.88 | 1.07 | -0.83 | **-1.54** | -1.27 | 0.05 | SP8 |
| sub21 | nTrans | 0.54 | -1.39 | **-2.40** | -0.17 | 0.55 | 0.61 | 0.61 | 0.59 | 0.52 | 0.54 | SP3 |
| sub22 | early-Trans | 0.64 | **-1.83** | -1.50 | -1.09 | 0.14 | 0.61 | 0.97 | 0.90 | 0.54 | 0.63 | SP2 |
| sub23 | nTrans | 0.37 | **-2.12** | -1.19 | 0.88 | 1.06 | -0.39 | 0.13 | 1.29 | -0.36 | 0.33 | SP2 |
| sub24 | nTrans | 0.67 | 0.81 | 0.85 | 1.02 | -0.39 | -0.99 | **-1.64** | -1.58 | 0.58 | 0.66 | SP7 |
| sub25 | Trans | 0.13 | 0.19 | -0.86 | -1.24 | **-1.52** | 1.23 | 1.44 | 1.17 | -0.63 | 0.09 | SP5 |
| sub26 | early-Trans | 0.78 | **-1.90** | -1.17 | -0.68 | -0.68 | 0.00 | 1.08 | 0.72 | 1.10 | 0.77 | SP2 |
| sub27 | late-Trans | 1.02 | 0.31 | 0.82 | 1.05 | -0.01 | -1.04 | -1.57 | **-1.60** | 0.01 | 1.00 | SP8 |
| sub28 | Trans | -0.02 | 0.93 | 0.29 | 0.65 | -1.22 | **-2.44** | 0.53 | 0.78 | 0.52 | -0.02 | SP6 |
| sub29 | nTrans | 0.00 | -1.60 | 1.30 | 0.84 | **-1.97** | -0.29 | 0.42 | 0.81 | 0.52 | -0.03 | SP5 |
| sub30 | nTrans | 0.82 | **-1.85** | -1.59 | -0.93 | 0.20 | 0.58 | 0.32 | 0.61 | 1.01 | 0.81 | SP2 |
| sub31 | late-Trans | 0.86 | 1.10 | 0.77 | 0.70 | -0.17 | -0.20 | -0.69 | -1.12 | **-2.11** | 0.85 | SP9 |
| sub32 | Trans | 0.78 | 1.01 | -0.12 | -0.85 | -1.58 | **-1.67** | 0.08 | 0.32 | 1.27 | 0.77 | SP6 |
| sub33 | Trans | 0.90 | 0.15 | **-2.18** | -1.33 | -0.21 | -0.24 | 0.19 | 0.72 | 1.09 | 0.90 | SP3 |
| sub34 | nTrans | 0.68 | 0.87 | 0.94 | -0.54 | 0.27 | 1.00 | -0.54 | -1.49 | **-1.88** | 0.69 | SP9 |
| sub35 | late-Trans | 0.21 | -0.77 | -0.86 | 1.04 | 1.05 | 0.82 | 0.92 | -0.45 | **-2.16** | 0.20 | SP9 |
| sub36 | Trans | -0.08 | 0.40 | **-2.58** | -0.82 | 0.16 | 0.78 | 0.43 | 0.84 | 0.97 | -0.11 | SP3 |
| sub37 | late-Trans | 0.72 | 1.08 | 0.82 | 0.68 | 0.57 | -0.45 | -1.62 | **-1.70** | -0.81 | 0.71 | SP8 |
| sub38 | early-Trans | **-1.66** | 0.64 | 0.72 | 0.76 | 0.76 | 0.42 | 0.64 | 0.58 | -1.17 | -1.69 | SP1 |
| sub39 | late-Trans | 1.49 | 1.14 | -0.24 | 0.09 | -0.66 | -0.51 | -0.21 | **-1.50** | -1.10 | 1.49 | SP8 |
| sub40 | Trans | 0.84 | -0.59 | **-2.34** | -0.89 | -0.19 | 0.01 | 0.45 | 0.92 | 0.96 | 0.84 | SP3 |
| sub41 | nTrans | -0.39 | 0.78 | 0.77 | 0.21 | 0.32 | 0.51 | 0.79 | 0.17 | **-2.72** | -0.45 | SP9 |
| sub42 | Trans | 1.26 | 0.93 | 0.71 | 0.24 | -0.05 | -0.70 | **-1.72** | -1.19 | -0.74 | 1.26 | SP7 |
| sub43 | nTrans | 0.78 | -1.66 | **-1.67** | -0.86 | -0.35 | 0.64 | 0.47 | 1.05 | 0.81 | 0.77 | SP3 |
| sub44 | Trans | 0.43 | 0.02 | **-2.08** | 0.43 | 1.34 | 1.17 | -0.61 | -1.26 | 0.17 | 0.38 | SP3 |
| sub45 | nTrans | **-1.56** | 1.43 | 1.06 | 0.58 | -0.77 | -0.09 | 0.68 | -0.38 | 0.62 | -1.57 | SP1 |
| sub46 | Trans | 0.12 | 1.47 | -1.10 | **-1.94** | -0.20 | -0.21 | 0.64 | 1.47 | -0.40 | 0.15 | SP4 |
| sub47 | nTrans | -1.04 | 0.88 | 0.19 | 0.56 | -1.18 | 1.16 | 0.96 | **-1.38** | 0.98 | -1.13 | SP8 |
| sub48 | nTrans | -1.26 | -0.80 | 0.82 | 0.84 | 1.31 | **-1.27** | 0.52 | 0.88 | 0.33 | -1.38 | SP6 |
| sub49 | early-Trans | 0.56 | **-1.59** | -0.11 | 1.39 | 0.91 | -0.63 | 0.98 | -0.48 | -1.58 | 0.56 | SP2 |
| sub50 | nTrans | -0.05 | -1.29 | **-1.66** | 0.83 | 1.62 | 0.71 | -0.96 | 0.01 | 0.91 | -0.13 | SP3 |
| sub51 | nTrans | **-1.37** | 1.43 | 1.17 | -0.38 | 0.50 | 1.02 | 0.25 | -1.11 | -0.12 | -1.40 | SP1 |
| sub52 | Trans | -0.43 | -1.04 | 1.45 | -1.13 | **-1.40** | 0.72 | 1.16 | 0.03 | 1.13 | -0.49 | SP5 |

*Continued on next page*

**Supplementary Table 3**: **Model-based switchpoint estimate for participants in Exp. 2.** Continued.

| Subject | Group | SP1 | SP2 | SP3 | SP4 | SP5 | SP6 | SP7 | SP8 | SP9 | No SP | Best SP |
|---|---|---|---|---|---|---|---|---|---|---|---|---|
| sub53 | Trans | 0.87 | 0.53 | 0.39 | -0.73 | -1.68 | **-1.85** | 0.28 | 0.91 | 0.24 | 1.03 | SP6 |
| sub54 | Trans | 0.95 | -1.42 | **-1.74** | -0.86 | -0.58 | 0.31 | 0.64 | 0.84 | 0.91 | 0.95 | SP3 |
| sub55 | Trans | 1.23 | -0.13 | -0.09 | 0.01 | -0.25 | **-2.13** | -0.43 | -0.72 | 1.29 | 1.23 | SP6 |
| sub56 | Trans | 1.05 | -1.23 | **-1.82** | 0.60 | 0.77 | -0.70 | -0.69 | -0.07 | 1.02 | 1.06 | SP3 |
| sub57 | nTrans | -0.82 | 1.70 | 1.21 | 0.64 | -0.39 | **-1.56** | -0.60 | -0.27 | 0.93 | -0.83 | SP6 |
| sub58 | late-Trans | -0.39 | 0.78 | 0.80 | 0.64 | 1.02 | 0.10 | 0.52 | **-2.53** | -0.55 | -0.40 | SP8 |
| sub59 | Trans | 0.07 | 1.03 | 0.88 | 1.16 | 0.05 | -0.21 | **-2.53** | -0.60 | 0.06 | 0.10 | SP7 |
| sub60 | early-Trans | **-1.31** | -0.21 | -0.83 | 1.40 | 1.24 | 1.30 | -0.30 | 0.42 | -0.33 | -1.40 | SP1 |
| sub61 | Trans | 0.52 | 0.69 | -0.41 | **-2.72** | -0.38 | 0.16 | 0.44 | 0.17 | 1.02 | 0.51 | SP4 |
| sub62 | nTrans | 0.20 | -1.48 | **-2.31** | -0.02 | 0.63 | 0.84 | 0.56 | 0.71 | 0.69 | 0.19 | SP3 |
| sub63 | nTrans | 0.32 | **-2.97** | 0.42 | 0.10 | 0.49 | 0.44 | 0.28 | 0.11 | 0.50 | 0.32 | SP2 |
| sub64 | nTrans | -0.18 | 0.90 | 0.96 | -0.32 | 0.88 | -0.93 | 0.50 | 0.76 | **-2.38** | -0.19 | SP9 |
| sub65 | Trans | 0.23 | 0.91 | 0.90 | 0.89 | -1.31 | **-2.17** | -0.46 | 0.91 | -0.13 | 0.23 | SP6 |
| sub66 | nTrans | 0.32 | 0.84 | -0.97 | **-2.58** | 0.01 | 0.72 | 0.89 | -0.01 | 0.46 | 0.32 | SP4 |
| sub67 | nTrans | **-1.45** | -0.48 | 1.01 | 0.88 | 1.04 | 0.21 | -1.22 | 0.51 | 0.95 | -1.46 | SP1 |
| sub68 | nTrans | 0.10 | **-1.86** | -1.23 | 0.15 | 0.82 | 1.31 | 0.87 | 0.84 | -1.07 | 0.06 | SP2 |
| sub69 | Trans | 0.05 | -1.72 | **-2.14** | 0.59 | 0.68 | 0.72 | 0.55 | 0.58 | 0.69 | 0.02 | SP3 |
| sub70 | nTrans | 0.49 | -1.76 | **-1.99** | -0.13 | -0.14 | 0.50 | 0.86 | 0.86 | 0.83 | 0.49 | SP3 |
| sub71 | early-Trans | 0.38 | **-2.21** | -1.02 | -0.88 | 0.16 | 0.80 | 0.46 | 0.63 | 1.33 | 0.35 | SP2 |
| sub72 | Trans | 0.14 | -0.11 | **-2.68** | -0.59 | 0.65 | 0.77 | 0.82 | 0.84 | 0.02 | 0.13 | SP3 |
| sub73 | nTrans | -0.42 | 0.71 | **-2.57** | 0.76 | 0.85 | -0.03 | 0.46 | 0.96 | -0.33 | -0.41 | SP3 |
| sub74 | nTrans | -0.36 | 0.23 | 0.28 | **-2.54** | -0.56 | 0.87 | 0.73 | 0.69 | 1.03 | -0.36 | SP4 |
| sub75 | early-Trans | 0.38 | **-2.93** | -0.27 | 0.49 | 0.42 | 0.30 | 0.22 | 0.42 | 0.60 | 0.36 | SP2 |
| sub76 | early-Trans | -0.04 | **-2.12** | 1.29 | 0.64 | 0.76 | -0.67 | 0.08 | -1.03 | 1.14 | -0.05 | SP2 |
| sub77 | Trans | 1.14 | -0.47 | -0.93 | 0.31 | **-2.03** | -0.17 | 0.75 | 1.00 | -0.72 | 1.13 | SP5 |
| sub78 | early-Trans | 0.41 | **-2.63** | -1.07 | 0.10 | 0.45 | 0.68 | 0.69 | 0.41 | 0.55 | 0.42 | SP2 |
| sub79 | Trans | 0.98 | 0.80 | 0.57 | 0.34 | -1.07 | -1.32 | **-1.85** | -0.24 | 0.81 | 0.98 | SP7 |
| sub80 | Trans | 0.90 | 1.04 | 0.93 | 0.56 | -0.17 | -0.70 | **-2.01** | -1.25 | -0.18 | 0.89 | SP7 |
| sub81 | nTrans | 0.16 | 1.32 | 0.81 | -0.30 | 0.02 | -0.44 | 0.30 | 0.55 | **-2.62** | 0.17 | SP9 |
| sub82 | nTrans | -0.19 | 0.83 | 0.76 | 0.68 | 0.64 | 0.54 | **-2.67** | -0.56 | 0.17 | -0.21 | SP7 |
| sub83 | nTrans | **-1.48** | 0.86 | 1.06 | 0.95 | 0.69 | 0.48 | 0.00 | -1.43 | 0.33 | -1.47 | SP1 |
| sub84 | late-Trans | -0.94 | -0.51 | -0.34 | 0.76 | 1.46 | -0.02 | 0.61 | **-1.09** | 1.53 | -1.47 | SP8 |
| sub85 | nTrans | 0.70 | **-2.38** | -1.25 | 0.69 | 0.18 | 0.31 | -0.33 | 0.95 | 0.44 | 0.69 | SP2 |
| sub86 | nTrans | -0.51 | 1.06 | 0.33 | -1.47 | **-1.87** | 0.20 | 0.91 | 1.01 | 0.83 | -0.50 | SP5 |

*Note.* For each participant, reaction time differences between regular and control sequences (ΔRT) were fitted using candidate single-switchpoint models and a no-switchpoint model. In the candidate switchpoint models, each learning session was treated in turn as the possible switchpoint. Transition status was defined independently of model fitting: participants were classified as transition learners only if they showed rule-attributed AU and ΔAU greater than 0 in the postlearning awareness test; all others were classified as non-transition learners. For transition learners, the transition session was then estimated from the best-fitting switchpoint model. Participants with estimated transitions in the first two or last two sessions were excluded from subsequent analyses because of insufficient decoding data (grouped as early-Trans or late-Trans). Values are normalized WAIC scores across ten model fits within participants, with lower values indicating better model fit. SP1–SP9 denote switchpoint models with the switchpoint placed at sessions 1–9, respectively. No SP denotes the no-switchpoint model. Best SP indicates the best-fitting switchpoint model among SP1–SP9 for the retained transition learners. In each row, the lowest value among SP1-SP9 is shown in bold.

**Supplementary Table 4: DK-atlas summary of source-space searchlight clusters for transition learners versus non-transition learners in Exp. 3.**

| Condition | Hemisphere | DK region | Significant vertices | Peak value | MNI x | MNI y | MNI z |
|---|---|---|---|---|---|---|---|
| Pre | Left | middle temporal | 12 | 0.014 | -57.08 | -42.30 | -11.15 |
| Pre | Left | banks of the superior temporal sulcus | 2 | 0.013 | -56.94 | -46.06 | -1.63 |
| Pre | Right | supramarginal | 13 | 0.012 | 36.95 | -39.08 | 35.92 |
| Pre | Right | superior parietal | 3 | 0.012 | 33.55 | -37.69 | 38.07 |
| Pre | Right | inferior parietal | 2 | 0.007 | 45.72 | -45.66 | 36.14 |
| Post | Left | superior temporal | 17 | 0.010 | -62.41 | -14.74 | -0.24 |
| Post | Left | rostral middle frontal | 13 | 0.013 | -42.16 | 30.59 | 20.78 |
| Post | Left | middle temporal | 7 | 0.016 | -54.37 | -60.10 | -1.53 |
| Post | Left | transverse temporal | 2 | 0.007 | -53.50 | -18.59 | 2.81 |
| Post | Left | banks of the superior temporal sulcus | 1 | 0.011 | -57.72 | -43.03 | -4.52 |
| Post | Left | inferior temporal | 1 | 0.010 | -55.78 | -55.20 | -10.88 |
| Post | Right | inferior parietal | 41 | 0.015 | 50.42 | -56.98 | 13.76 |
| Post | Right | supramarginal | 27 | 0.017 | 46.31 | -30.45 | 38.70 |
| Post | Right | middle temporal | 20 | 0.013 | 52.38 | -53.82 | -4.57 |
| Post | Right | banks of the superior temporal sulcus | 15 | 0.010 | 50.00 | -46.86 | 6.41 |
| Post | Right | postcentral | 15 | 0.015 | 41.75 | -28.68 | 46.71 |
| Post | Right | pars triangularis | 14 | 0.014 | 47.39 | 30.28 | -0.55 |
| Post | Right | pars orbitalis | 11 | 0.017 | 41.60 | 42.43 | -12.66 |
| Post | Right | inferior temporal | 7 | 0.016 | 49.61 | -58.40 | -7.89 |
| Post | Right | lateral orbitofrontal | 4 | 0.011 | 30.99 | 40.20 | -9.73 |
| Post | Right | fusiform | 2 | 0.013 | 42.66 | -57.05 | -9.75 |
| Post | Right | lateral occipital | 1 | 0.008 | 42.46 | -77.39 | 6.81 |
| Post − pre | Left | insula | 24 | 0.016 | -35.60 | -3.27 | -5.49 |
| Post − pre | Left | superior frontal | 14 | 0.018 | -13.24 | -10.01 | 44.27 |
| Post − pre | Left | rostral middle frontal | 12 | 0.018 | -33.66 | 47.95 | 17.36 |
| Post − pre | Left | posterior cingulate | 11 | 0.017 | -5.14 | -3.55 | 30.35 |
| Post − pre | Left | unassigned | 9 | 0.016 | -4.18 | -23.66 | -14.48 |
| Post − pre | Left | precentral | 5 | 0.013 | -46.48 | -1.97 | 11.85 |
| Post − pre | Left | postcentral | 4 | 0.011 | -42.22 | -18.06 | 19.19 |
| Post − pre | Left | supramarginal | 4 | 0.016 | -37.94 | -26.98 | 22.13 |
| Post − pre | Left | lateral orbitofrontal | 1 | 0.012 | -27.27 | 18.92 | -6.43 |
| Post − pre | Right | superior frontal | 28 | 0.021 | 18.93 | -0.69 | 65.45 |
| Post − pre | Right | superior temporal | 26 | 0.017 | 53.12 | -31.30 | 1.85 |
| Post − pre | Right | middle temporal | 21 | 0.017 | 62.08 | -47.01 | -4.81 |
| Post − pre | Right | banks of the superior temporal sulcus | 20 | 0.019 | 45.02 | -41.37 | 2.36 |
| Post − pre | Right | inferior parietal | 17 | 0.016 | 48.59 | -56.40 | 9.13 |
| Post − pre | Right | pars orbitalis | 12 | 0.020 | 36.93 | 38.53 | -8.14 |
| Post − pre | Right | supramarginal | 10 | 0.014 | 35.78 | -32.62 | 15.28 |
| Post − pre | Right | insula | 9 | 0.016 | 32.27 | -24.20 | 17.99 |
| Post − pre | Right | pars triangularis | 7 | 0.017 | 42.29 | 39.33 | -0.69 |
| Post − pre | Right | rostral middle frontal | 7 | 0.012 | 39.63 | 50.45 | -1.91 |
| Post − pre | Right | cuneus | 6 | 0.014 | 5.05 | -83.56 | 10.10 |
| Post − pre | Right | inferior temporal | 5 | 0.015 | 53.86 | -55.66 | -7.26 |
| Post − pre | Right | paracentral | 5 | 0.012 | 7.58 | -38.82 | 59.71 |
| Post − pre | Right | postcentral | 4 | 0.011 | 8.53 | -39.25 | 74.49 |
| Post − pre | Right | transverse temporal | 4 | 0.014 | 42.38 | -23.40 | 7.68 |

*Continued on next page*

**Supplementary Table 4**: **DK-atlas summary of source-space searchlight clusters for transition learners versus non-transition learners in Exp. 3.** Continued.

| Condition | Hemisphere | DK region | Significant vertices | Peak value | MNI x | MNI y | MNI z |
|---|---|---|---|---|---|---|---|
| Post − pre | Right | lateral orbitofrontal | 2 | 0.014 | 32.49 | 35.08 | -6.73 |
| Post − pre | Right | pericalcarine | 2 | 0.010 | 8.52 | -75.69 | 14.21 |

*Note.* Cortical source vertices were assigned to Desikan–-Killiany (DK; FreeSurfer aparc) regions on fsaverage. For each condition, hemisphere, and DK region, the table reports the number of source vertices surviving cluster-based correction ($P < 0.05$) in the corresponding surface map, the peak (maximum) compression-rate difference within those vertices, and the MNI coordinates of the peak vertex. *Pre* and *Post* denote state-specific group-difference maps computed as transition minus non-transition learners within the pre- and post-transition states, respectively. *Post − pre* denotes the change in this group difference across states, computed as $(\text{transition} - \text{non-transition})_{\text{post}} - (\text{transition} - \text{non-transition})_{\text{pre}}$. "Unassigned" denotes significant vertices not assigned to a named DK region. Peak values are rounded to three decimal places and MNI coordinates to two decimal places.

**Supplementary Table 5: DK-atlas summary of source-space ROIs used for phase-transfer entropy analyses in Exp. 3.**

| Source map | Cluster | Hemisphere | DK region | Significant vertices | Peak value | MNI x | MNI y | MNI z |
|---|---|---|---|---|---|---|---|---|
| Pre | 1 | Left | **middle temporal** | **14 (53.8%)** | **0.015** | **-62.16** | **-38.20** | **-9.61** |
| Pre | 1 | Left | inferior temporal | 10 (38.5%) | 0.013 | -52.81 | -35.17 | -18.45 |
| Pre | 1 | Left | banks of the superior temporal sulcus | 2 (7.7%) | 0.011 | -56.94 | -46.06 | -1.63 |
| Post | 2 | Left | **rostral middle frontal** | **24 (49.0%)** | **0.013** | **-42.16** | **30.59** | **20.78** |
| Post | 2 | Left | caudal middle frontal | 16 (32.7%) | 0.012 | -34.17 | 13.95 | 25.13 |
| Post | 2 | Left | pars opercularis | 8 (16.3%) | 0.010 | -37.37 | 14.42 | 23.01 |
| Post | 2 | Left | precentral | 1 (2.0%) | 0.008 | -42.29 | -0.84 | 29.20 |
| Post | 3 | Left | **parahippocampal** | **10 (40.0%)** | 0.012 | -28.78 | -41.83 | -8.92 |
| Post | 3 | Left | lingual | 8 (32.0%) | 0.013 | -32.96 | -44.30 | -7.51 |
| Post | 3 | Left | fusiform | 6 (24.0%) | **0.014** | **-34.62** | **-46.94** | **-10.38** |
| Post | 3 | Left | entorhinal | 1 (4.0%) | 0.008 | -22.23 | -21.66 | -22.61 |
| Post | 4 | Right | **pars triangularis** | **27 (50.0%)** | 0.014 | 40.90 | 30.14 | 2.52 |
| Post | 4 | Right | pars orbitalis | 14 (25.9%) | **0.014** | **36.93** | **38.53** | **-8.14** |
| Post | 4 | Right | lateral orbitofrontal | 10 (18.5%) | 0.013 | 32.49 | 35.08 | -6.73 |
| Post | 4 | Right | insula | 2 (3.7%) | 0.010 | 31.15 | 21.51 | 1.74 |
| Post | 4 | Right | rostral middle frontal | 1 (1.9%) | 0.008 | 35.40 | 34.29 | 12.99 |
| Post | 5 | Right | **pars opercularis** | **14 (41.2%)** | 0.012 | 48.64 | 7.43 | 2.73 |
| Post | 5 | Right | precentral | 10 (29.4%) | **0.014** | **57.52** | **4.53** | **8.23** |
| Post | 5 | Right | rostral middle frontal | 9 (26.5%) | 0.013 | 45.00 | 28.48 | 23.92 |
| Post | 5 | Right | caudal middle frontal | 1 (2.9%) | 0.009 | 32.96 | 13.68 | 27.68 |
| Post | 6 | Right | **supramarginal** | **39 (37.1%)** | **0.017** | **44.86** | **-36.48** | **41.30** |
| Post | 6 | Right | postcentral | 30 (28.6%) | 0.017 | 41.75 | -28.68 | 46.71 |
| Post | 6 | Right | inferior parietal | 26 (24.8%) | 0.013 | 47.47 | -54.78 | 44.80 |
| Post | 6 | Right | superior parietal | 6 (5.7%) | 0.011 | 33.55 | -37.69 | 38.07 |
| Post | 6 | Right | precentral | 4 (3.8%) | 0.010 | 33.99 | -21.28 | 42.33 |
| Post | 7 | Right | **parahippocampal** | **17 (41.5%)** | 0.012 | 34.39 | -32.79 | -15.44 |
| Post | 7 | Right | lingual | 13 (31.7%) | 0.014 | 27.94 | -47.07 | -6.01 |
| Post | 7 | Right | fusiform | 10 (24.4%) | **0.017** | **32.41** | **-57.33** | **-8.54** |
| Post | 7 | Right | isthmus cingulate | 1 (2.4%) | 0.007 | 15.05 | -40.34 | -2.26 |
| Post | 8 | Right | **inferior parietal** | **43 (19.7%)** | 0.013 | 50.42 | -56.98 | 13.76 |
| Post | 8 | Right | fusiform | 41 (18.8%) | 0.018 | 35.73 | -57.90 | -16.02 |
| Post | 8 | Right | supramarginal | 31 (14.2%) | 0.013 | 49.63 | -37.86 | 23.01 |
| Post | 8 | Right | inferior temporal | 20 (9.2%) | **0.019** | **49.61** | **-58.40** | **-7.89** |
| Post | 8 | Right | banks of the superior temporal sulcus | 19 (8.7%) | 0.010 | 54.65 | -39.79 | 12.82 |
| Post | 8 | Right | lateral occipital | 19 (8.7%) | 0.013 | 42.11 | -69.23 | -5.45 |
| Post | 8 | Right | middle temporal | 18 (8.3%) | 0.014 | 52.38 | -53.82 | -4.57 |
| Post | 8 | Right | superior temporal | 16 (7.3%) | 0.011 | 53.12 | -31.30 | 1.85 |
| Post | 8 | Right | insula | 8 (3.7%) | 0.014 | 34.40 | -23.80 | 20.65 |
| Post | 8 | Right | parahippocampal | 3 (1.4%) | 0.011 | 36.61 | -29.93 | -17.76 |

*Note.* Cortical source vertices in each phase-transfer entropy (PTE) ROI were assigned to Desikan–Killiany (DK; FreeSurfer aparc) regions on fsaverage. Cluster identifiers follow the node numbers used in the finalized seed-ROI PTE surface legend. *Pre* and *Post* denote the transition learners pre- and post-transition source maps from which the ROIs were selected. For each legend-defined cluster and DK region, the table reports the number of selected DK-assigned vertices and, in parentheses, the percentage of the DK-assigned cluster total. Within each cluster, the dominating DK-region row and its vertex count are shown in bold. Bold peak values and MNI coordinates mark the DK-region row containing the maximum compression-rate value within that cluster. Peak values are rounded to three decimal places and MNI coordinates to two decimal places.

### Supplementary Figures

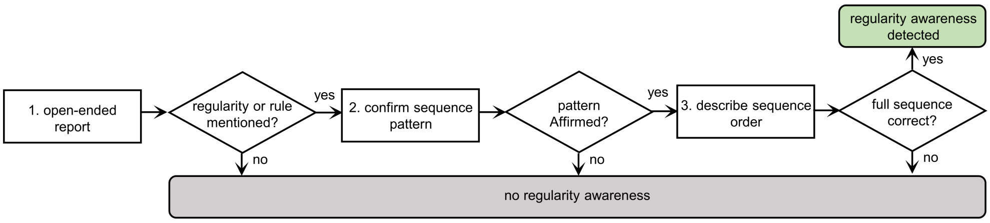

**Supplementary Fig. 1**: **Protocol of session-wise awareness test.** At the end of each learning session, participants completed a verbal awareness test designed to detect spontaneous awareness without giving explicit hints about the sequence rule. In Step 1, the experimenter posed an open-ended question: "Please describe any feelings about the previous task session. There are no right or wrong answers. Any feelings are valuable and should be reported." Participants continued reporting until they explicitly indicated that they had reported everything they noticed or felt. Reports referring only to subjective emotions, picture content, the experimental environment, or other sequence-irrelevant impressions, without mentioning order, sequence, rules, or patterns, were classified as no regularity awareness, and the assessment for the current session was terminated. Reports mentioning order, sequence, rules, or patterns in the image stimuli or button presses were treated as potential regularity awareness and led to Step 2. In Step 2, the experimenter asked a probing confirmation question, using the participant's own terminology where possible to avoid providing additional hints, for example, "Do you mean there is a sequence regularity among these images?" A negative response terminated the assessment and was coded as no regularity awareness. An affirmative response led to Step 3, in which participants were asked to describe the regularity they had noticed. Descriptions could refer to the complete sequence, partial transitions, or specific positions, such as a face appearing first. Participants were classified as having regularity awareness only when they correctly reported the complete trained sequence order, for example, face → tool → animal for type I learners. The first session in which this criterion was met was defined as the transition session, marking the shift from the implicit to the explicit state of sequence-regularity learning. All other response paths were coded as no regularity awareness before participants continued to the next session.

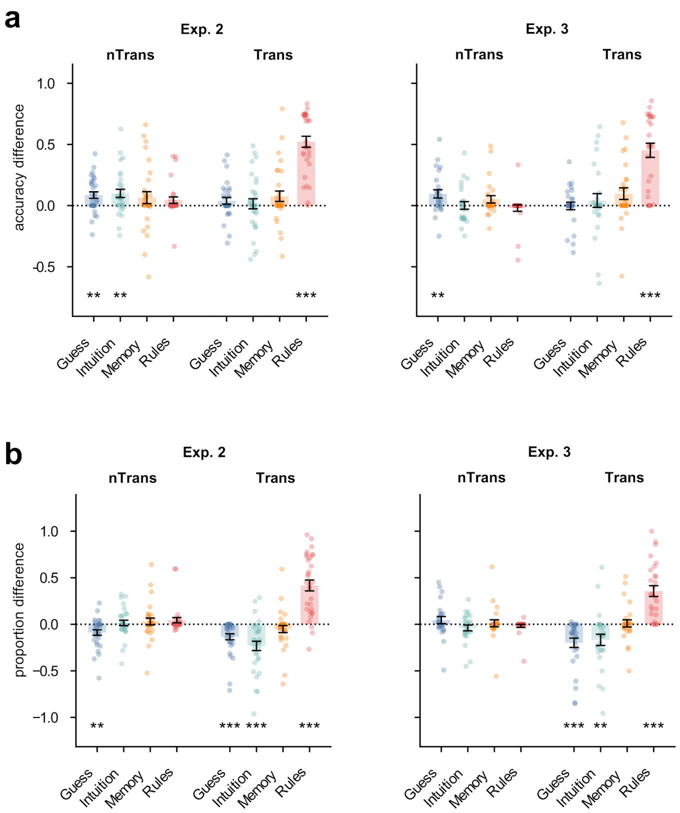

**Supplementary Fig. 2**: **Accuracy and attribution proportion in the postlearning awareness test in Exps. 2 and 3. a**, Accuracy difference was calculated as accuracy for the trained regular sequence minus accuracy for the alternative regular sequence and is plotted separately for non-transition and transition learners. To obtain defined accuracy estimates when a given attribution was not selected, accuracy was computed with a correction: $(N_\mathrm{c} + 0.5)$ / $(N + 1)$, where $N_\mathrm{c}$ denotes the number of correct-response trials assigned to a given attribution and $N$ denotes the number of trials assigned to that attribution. Transition learners showed a predominant accuracy advantage for "Rules" attributions (one-sample $t$ tests against zero, $t$s $\geq 7.80$, $P$s $< 0.001$, $d_z$s $\geq 1.50$). Non-transition learners showed accuracy advantages primarily for "Guess" attributions ($t$s $\geq 2.82$, $P$s $\leq 0.010$, $d_z$s $\geq 0.56$) across Exps. 2 and 3, and additionally for "Intuition" in Exp. 2 ($t = 2.98$, $P = 0.006$, $d_z = 0.54$). Points show individual learners; error bars denote s.e.m. **b**, Proportion difference was computed as the fraction of correct responses assigned to each attribution for the trained regular sequence minus the corresponding fraction for the alternative regular sequence. Transition learners preferentially attributed correct responses to "Rules" (one-sample $t$ tests against zero, $t$s $\geq 6.14$, $P$s $< 0.001$, $d_z$s $\geq 1.18$) and less to "Guess" or "Intuition" ($|t|$s $\geq 2.77$, $P$s $\leq 0.010$, $|d_z|$s $\geq 0.53$). In non-transition learners, no such preferential attribution pattern was observed in Exps. 2 and 3. Plot conventions are as in panel **a**.

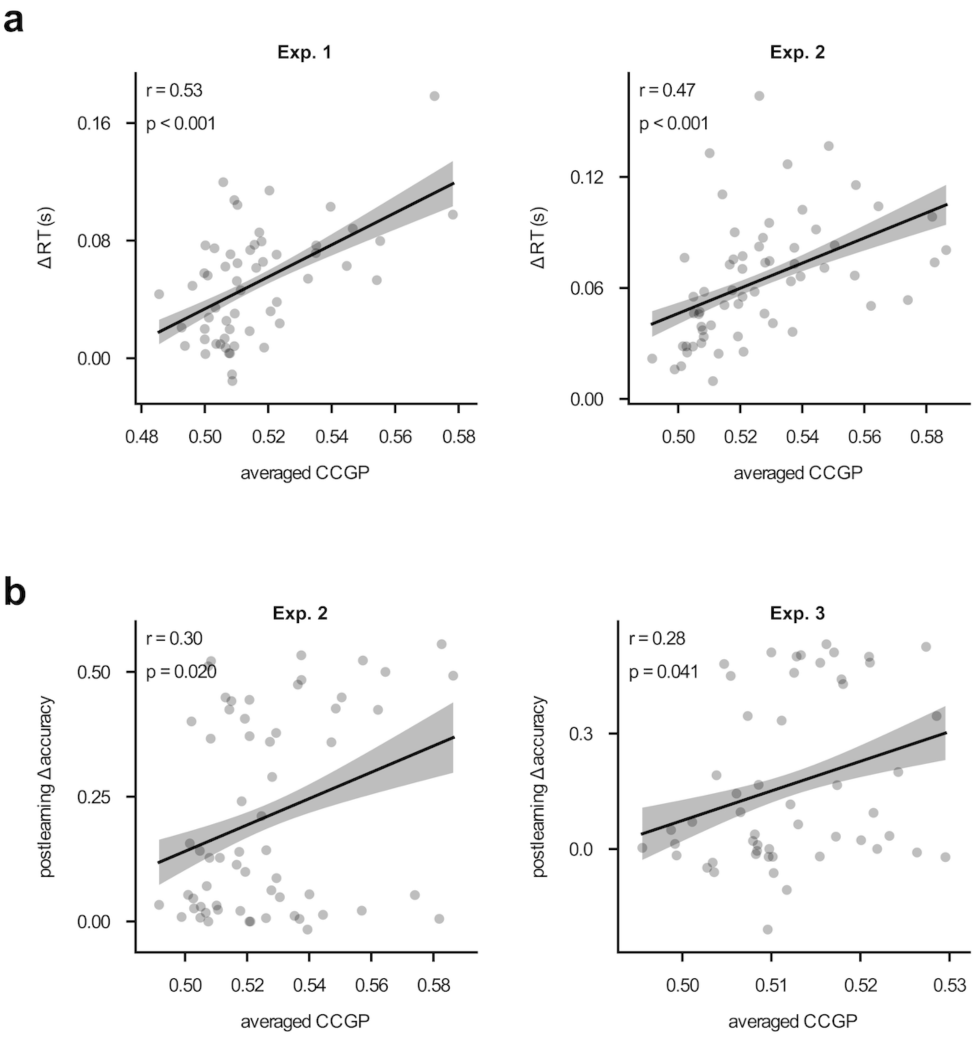

**Supplementary Fig. 3**: **Mean CCGP and its behavioural relevance.** Correlations between mean CCGP and behavioural indices of RT (**a**) in Exps. 1 and 2 and postlearning accuracy (**b**) in Exps. 2 and 3. CCGP was averaged within the significant time window identified across groups (transition vs. non-transition) and states (pre- vs. post-transition; i.e., the cluster-corrected window in Fig. 2b). $\Delta$RT ($RT_{control} - RT_{regular}$) was averaged across sessions in Exps. 1 and 2. Postlearning discrimination accuracy in Exps. 2 and 3 was quantified as $\Delta$Accuracy (trained regular accuracy − alternative regular accuracy; collapsed across attributions). Each dot denotes an individual learner; solid lines show Pearson linear fits and shaded bands denote s.e.m.

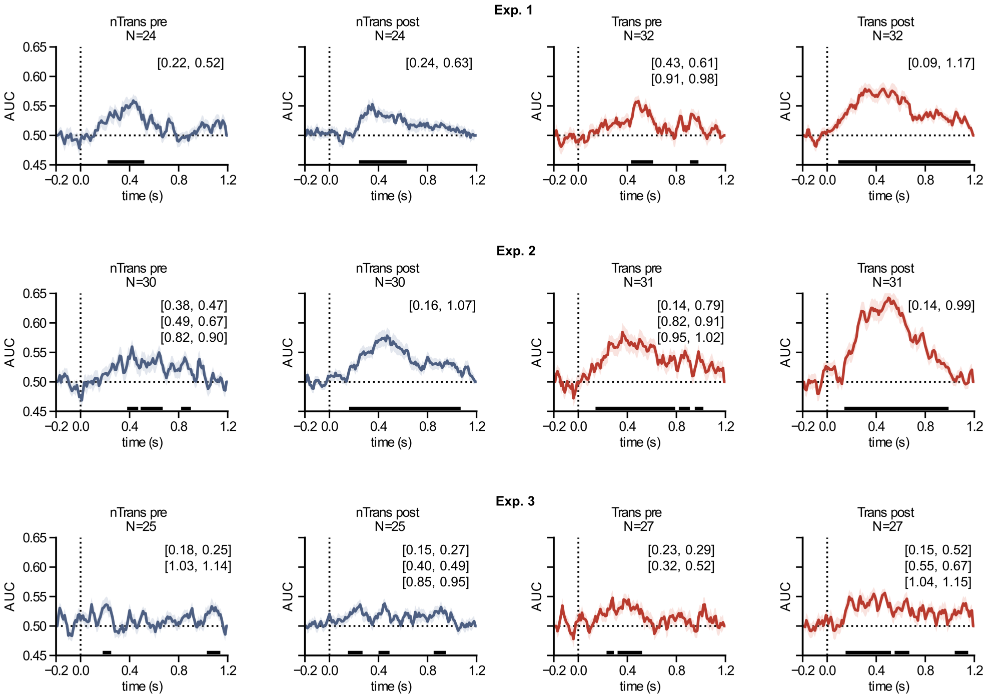

**Supplementary Fig. 4**: **Within-element decoding across participant groups and awareness states.** Time-resolved within-element decoding performance for Exps. 1–3, plotted separately for pre-transition and post-transition states in transition learners (Trans) and for matched early and late states in non-transition learners (nTrans). Decoders classified regular versus control trials within the same trained second- or third-position context, with regular and control trials matched for both sequence position and stimulus category. This analysis used the same trained decoders as the CCGP analysis but tested them on the same element rather than across elements. Five-fold cross-validation was used for classifier training and testing. AUC time courses were obtained by combining classifier evidence across the two trained contexts. Shading denotes the s.e.m. across participants. The horizontal dotted line indicates chance performance (AUC = 0.5), and the vertical dotted line (Time 0) indicates stimulus onset. Black horizontal bars mark significant above-chance time clusters, determined using a 1,000-permutation cluster-corrected test with $P < 0.05$; corresponding cluster intervals are shown in each panel.

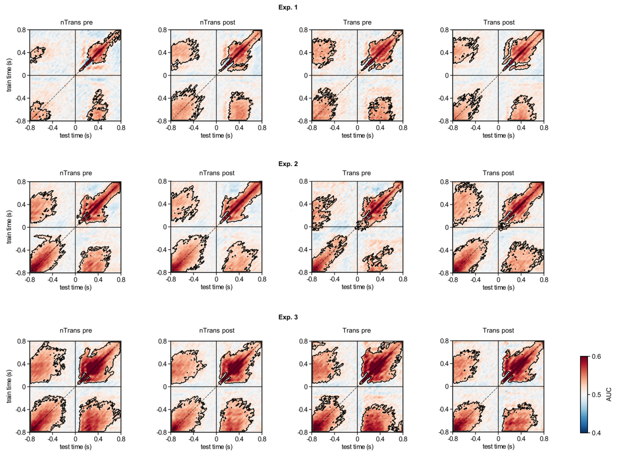

**Supplementary Fig. 5**: **Upcoming predictable stimulus categories were decodable before stimulus onset in both implicit and explicit states.** Time-generalization matrices show decoding of the successive predictable stimulus categories in Exps. 1–3, plotted separately for pre-transition and post-transition states in transition learners (Trans) and for matched early and late states in non-transition learners (nTrans). Decoders classified the second versus third positions of regular trials, corresponding to the two stimulus categories predictably specified by the trained sequence context. Classifiers were trained at each time point and tested at every other time point, yielding AUC values as a function of train time and test time relative to stimulus onset. Negative times indicate the inter-trial interval (ITI) before stimulus onset, and positive times indicate post-stimulus activity. The dashed diagonal marks within-time decoding, and the vertical and horizontal solid lines mark stimulus onset. Black contours indicate significant above-chance clusters, determined using a cluster-corrected test against chance performance (AUC = 0.5) with $P < 0.05$. Significant clusters in the upper-left and lower-right quadrants indicate bidirectional cross-temporal generalization between pre-stimulus ITI activity and post-stimulus evoked activity. Thus, classifiers trained on post-stimulus responses decoded the predictable category during the preceding ITI, whereas classifiers trained on ITI activity decoded the subsequent post-stimulus response. This pattern was observed across awareness states and learner groups, indicating that learned sequence context pre-activated the upcoming stimulus category even before sensory input arrived. This ITI decoding therefore captures stimulus-level perceptual prediction within regular trials. It is distinct from cross-condition generalization performance and representational-geometry analyses, which test whether regularity information generalizes across different elements or ordinal positions. Together, these results suggest that sequence learning can support anticipatory category-specific representations during the ITI, while partly separable analyses assess the more abstract, element-general organization of sequence representations.

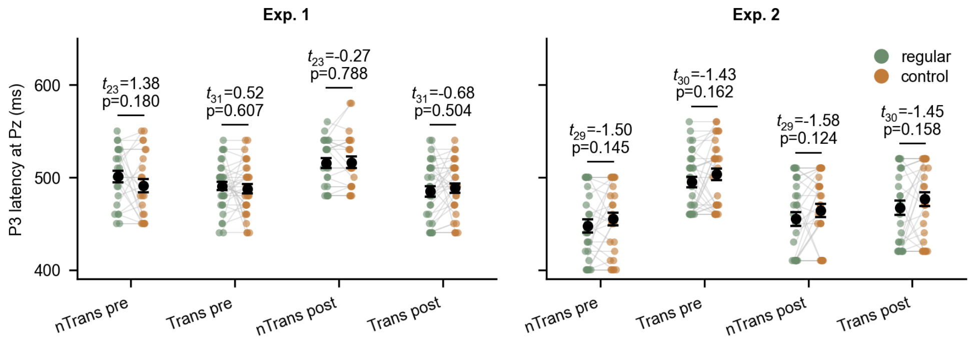

**Supplementary Fig. 6**: **Participant-wise P3 latency did not differ between regular and control trials in Exps. 1 and 2.** P3 peak latency was quantified at the Pz electrode for each participant, separately for regular and control trials, learner group and pre-/post-transition state. For each experiment, group and state, latency was estimated within the independently defined group-level 100-ms P3 peak window used for the ERP analyses. Within that window, we identified the time point of the maximum positive Pz amplitude separately for regular and control trials. Regular and control P3 latencies were then compared using two-tailed paired $t$ tests within each experiment, group and state. Dots show individual participants, grey lines connect paired regular and control estimates from the same participant, and black markers with error bars denote the mean $\pm$ s.e.m. Across Exps. 1 and 2, no regular–control P3 latency difference was significant in any learner group or pre-/post-transition state (paired $t$ tests, $|t|$s $\leq 1.58$, $P$s $\geq 0.124$, $|d_z|$s $\leq 0.29$). These results indicate that the P3 amplitude effect in transition learners and the awareness-predictive P3-window compression effect were not explained by P3 latency.
